# Variations of intracellular density during the cell cycle arise from tip-growth regulation in fission yeast

**DOI:** 10.1101/2020.10.21.349696

**Authors:** Pascal D. Odermatt, Teemu P. Miettinen, Joon Ho Kang, Emrah Bostan, Scott Manalis, Kerwyn Casey Huang, Fred Chang

## Abstract

Intracellular density impacts the physical nature of the cytoplasm and can globally affect cellular processes, yet density regulation remains poorly understood. Here, using a new quantitative phase imaging method, we determined that dry-mass density varies during the cell cycle in fission yeast. Density decreased during G2, increased in mitosis and cytokinesis, and rapidly dropped at cell birth. These density variations were explained by a constant rate of biomass synthesis, coupled to slowdown of volume growth during cell division and rapid expansion post-cytokinesis. Arrest at specific cell-cycle stages led to continued increases or decreases in density. Spatially heterogeneous patterns of density suggested links between density regulation and tip growth, and septum bending away from higher-density daughters linked density to intracellular osmotic pressure. Our results demonstrate that systematic density variations during the cell cycle are predominantly due to modulation of volume expansion, and reveal functional consequences of density gradients and cell-cycle arrests.

## Introduction

Intracellular density, a cumulative measure of the concentrations of all cellular components, is an important parameter that globally affects cellular function: density affects the concentration and activities of biomolecules, and can impact biophysical properties of the cytoplasm such as macromolecular crowding, diffusion, mechanical stiffness, and phase transitions (1–4). Although it is often assumed that density must be maintained at a particular level to optimize fitness, there is a growing appreciation that intracellular density often varies across physiological conditions. For example, substantial shifts in density and/or crowding have been detected in development, aging, and disease states (3, 5). Even normal cell-cycle progression can involve changes in density; in cultured mammalian cells, volume increases by 10-30% during mitosis (6, 7), which likely dilutes the cytoplasm prior to an increase in density during cytokinesis.

Despite this biological centrality, the homeostatic mechanisms maintaining cellular density remain poorly understood. Over the course of a typical cell cycle, cells both double their volume and duplicate all cellular contents. A critical unresolved question is how this growth in cell volume (increase in cell size) and biosynthesis are coordinated. In walled cells, the rate of volume growth is dictated largely by cell-wall synthesis and turgor pressure (8). In principle, feedback mechanisms could tightly couple biosynthesis with wall expansion. However, recent studies have demonstrated that it is possible to decouple biosynthesis from volume growth. For instance, budding yeast cells that are arrested in G1 phase grow to very large sizes and exhibit dilution of the cytoplasm accompanied by decreased protein synthesis and growth rate (9). Conversely, temporary inhibition of volume growth using osmotic oscillations or inhibition of secretion leads to increased density accompanied by a subsequent dramatic increase in volume growth rate in fission yeast (10).

Numerous methods have been developed to measure aspects of biomass and intracellular density and crowding in living cells (2, 3, 11). Suspended microchannel resonators (SMRs) infer buoyant cell mass from changes in the frequency of a resonating cantilever as a single cell passes through an embedded microchannel (12). As an alternative to SMRs, quantitative phase imaging (QPI) is a well-established optical technique for extracting dry-mass measurements from changes in the refractive index (13). The interpretation of phase shifts depends on the similarity in the refractive indices of major cellular components such as proteins, lipids, and nucleic acids (11). Previous studies have used phase gratings or holography to generate phase-shift maps that can be used to quantify intracellular density; these approaches require specialized equipment (14). Precise measurement of density throughout the cell cycle requires non-invasive, long-term quantification of single cells at high spatial and temporal resolution.

Here, we developed a new QPI method for measuring the intracellular density of fission yeast cells. This label-free method is based on the analysis of *z*-stacks of brightfield images (15), thus having the advantage of not requiring a specialized phase objective or holographic system. The fission yeast *Schizosaccharomyces pombe* is a leading cell-cycle model, as these rod-shaped cells have a highly regular shape, size, and cell cycle conducive to quantitative analyses. Using QPI to track the density of individual cells as they grew and divided within a microfluidic chamber, we determined that wild-type fission yeast cells exhibit characteristic density changes during the cell cycle, in which density falls during G2 phase and increases during mitosis and cytokinesis. These density variations arise from differences in the relative rates of volume and mass growth; while mass grows exponentially throughout the cell cycle, volume growth varies dependent on cell cycle stage/phase. Perturbations to cell-cycle progression and/or growth exacerbated density changes. We further observed gradients in density within single cells and found that intracellular density variations were correlated with tip growth and intracellular pressure. Our findings illustrate a general mechanism by which density is regulated through controlling the relative rates of volume growth and biosynthesis.

## Results

### QPI enables high-resolution measurements of intracellular density in growing cells

To measure intracellular density using a standard wide-field microscope, we developed a version of QPI in which the phase shift is retrieved computationally from a *z-*stack of brightfield images (15). This label-free approach takes advantage of the relationship between the intracellular concentration of biomolecules and the refractive index of the cell interior (11), which can be computed from the light intensity profile along the *z*-direction using the transport-of-intensity equation (Figure 1A, Methods). To calibrate phase shifts with absolute concentrations (dry-mass/volume), we measured the phase shifts within cells grown in media containing a range of concentrations of a calibration standard (bovine serum albumin, BSA) (Methods); these measurements showed a linear relationship that can be used to extrapolate the intracellular density of cells (Figure 1B). This method provides pixel-scale measurements of density in living cells and can easily be applied during time-lapse imaging with sub-minute time resolution on most wide-field microscopes.

**Figure 1:**
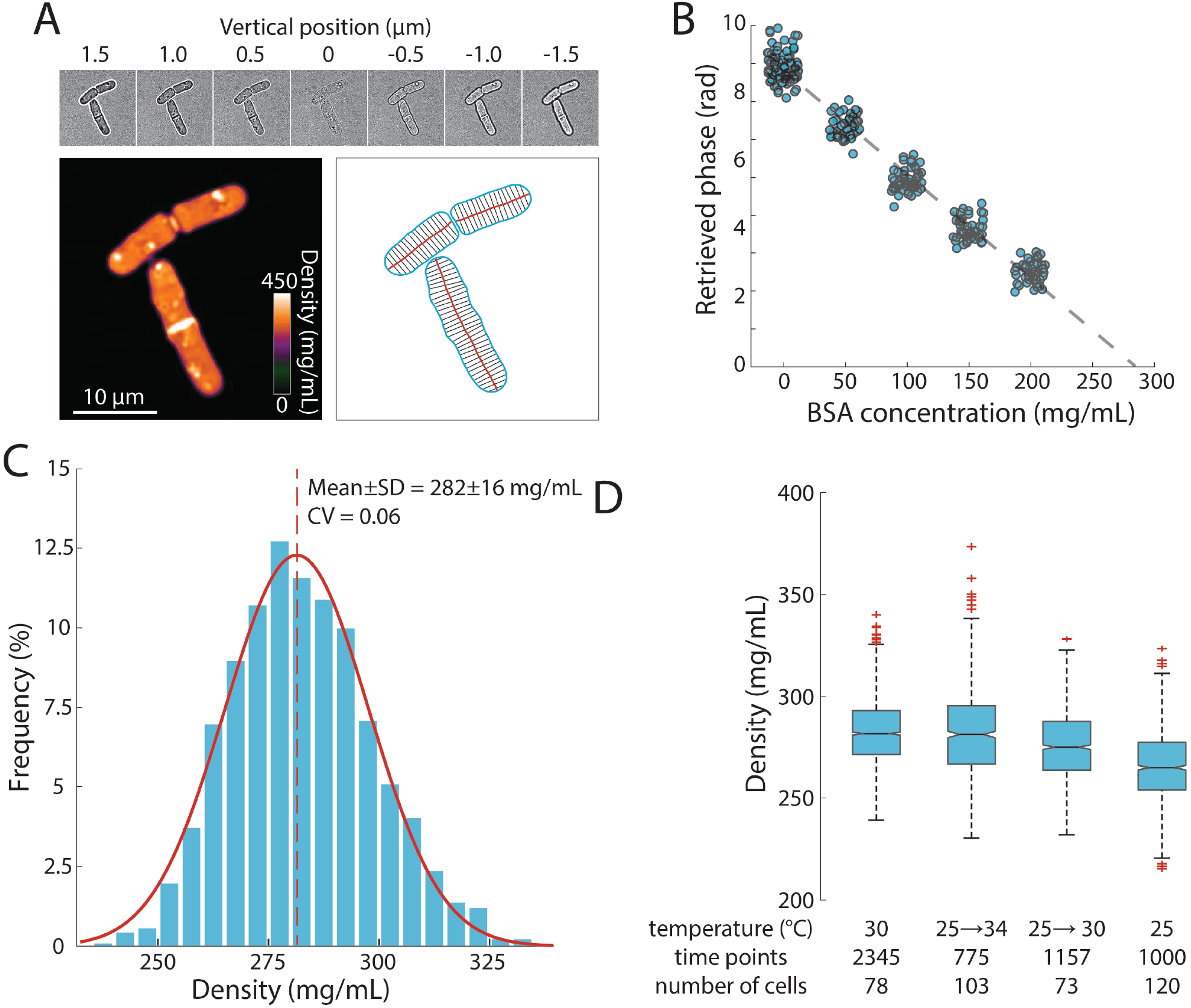
Precise measurement of intracellular density using QPI based on a *z*-stack of brightfield images. A) QPI method for computing cytoplasmic density from brightfield images. A *z*-stack of brightfield images of fission yeast cells ±1.5 μm around the focal position (top) were computationally analyzed by solving the transport-of-intensity equation (15) to retrieve pixel-by-pixel phase-shift maps (bottom left). Cellular dimensions were determined via segmentation and skeletonization (bottom right). B) QPI phase shifts were calibrated by imaging cells in media supplemented with a range of concentrations of BSA. The retrieved phase shift is linearly related to concentration (dashed line is the linear best fit). C) Histogram of dry-mass density measurements of exponential-phase fission yeast cells grown at 30 °C in YE5S medium. A Gaussian fit (red) yielded a mean (dashed line) density of 282±16 mg/mL (*n*=2345 time points, 78 cells). SD, standard deviation. D) Average cell density varied by less than 10% across temperatures and temperature shifts. Edges of the boxes indicate 25^th^ and 75^th^ percentiles, centerline is the median, and whiskers indicate extreme points not considered outliers. In experiments in which the temperature was shifted up, cells were initially grown at 25 °C and then moved to the microscope pre-heated to the indicated temperature for long-term imaging.

Using this methodology, we determined that the mean dry-mass density of an asynchronous population of wild-type *S. pombe* cells growing at 30 °C in rich YE5S medium was 282±16 mg/mL (Figure 1C). The distribution of densities was remarkably narrow, with a coefficient of variation (CV) of 0.06, despite variability in cell size, cell-cycle stage, and various sources of intracellular heterogeneities such as lipid droplets (Figure S1) and cell-wall septa (both of which were regions of high signal, Figure 1A). For cells within the same cell-cycle stage, the distribution of densities was even more narrow (CV<0.05, Figure S1B). The nucleus was not distinguishable in most phase-shift maps, indicating similar density as the cytoplasm (Figure 1A). A similar distribution of densities was observed in cells grown at 25 °C and after temperature shifts (Figure 1D). Together, these results suggest that dry-mass density is robustly maintained, and demonstrate that this QPI approach can precisely measure absolute dry-mass density in living cells with high temporal and spatial resolution.

### Intracellular density follows a characteristic trajectory during the *S. pombe* cell cycle

To determine whether intracellular density changes over the course of the fission yeast cell cycle, we imaged proliferating cells in time-lapse using QPI in a microfluidic device under constant flow of growth medium (Methods; Figure 2A, Movie S1). Density maps were segmented to extract cellular dimensions, from which volume was computed (Methods; Figure 1A). Total dry mass of each cell was computed from volume and mean density measurements. We imaged cells throughout their entire cell cycle, and then aligned the computed data from each cell by relative cell-cycle progression, from cell birth (first detectable physical separation between daughter cells) until just before cell-cell separation at the end of the cell cycle. As observed previously (16), cells exhibited steady tip growth in interphase (mostly G2 phase), and then volume growth slowed or halted during mitosis and cytokinesis (defined here as the period starting from septum formation and ending at daughter-cell separation; Figure 2B).

**Figure 2:**
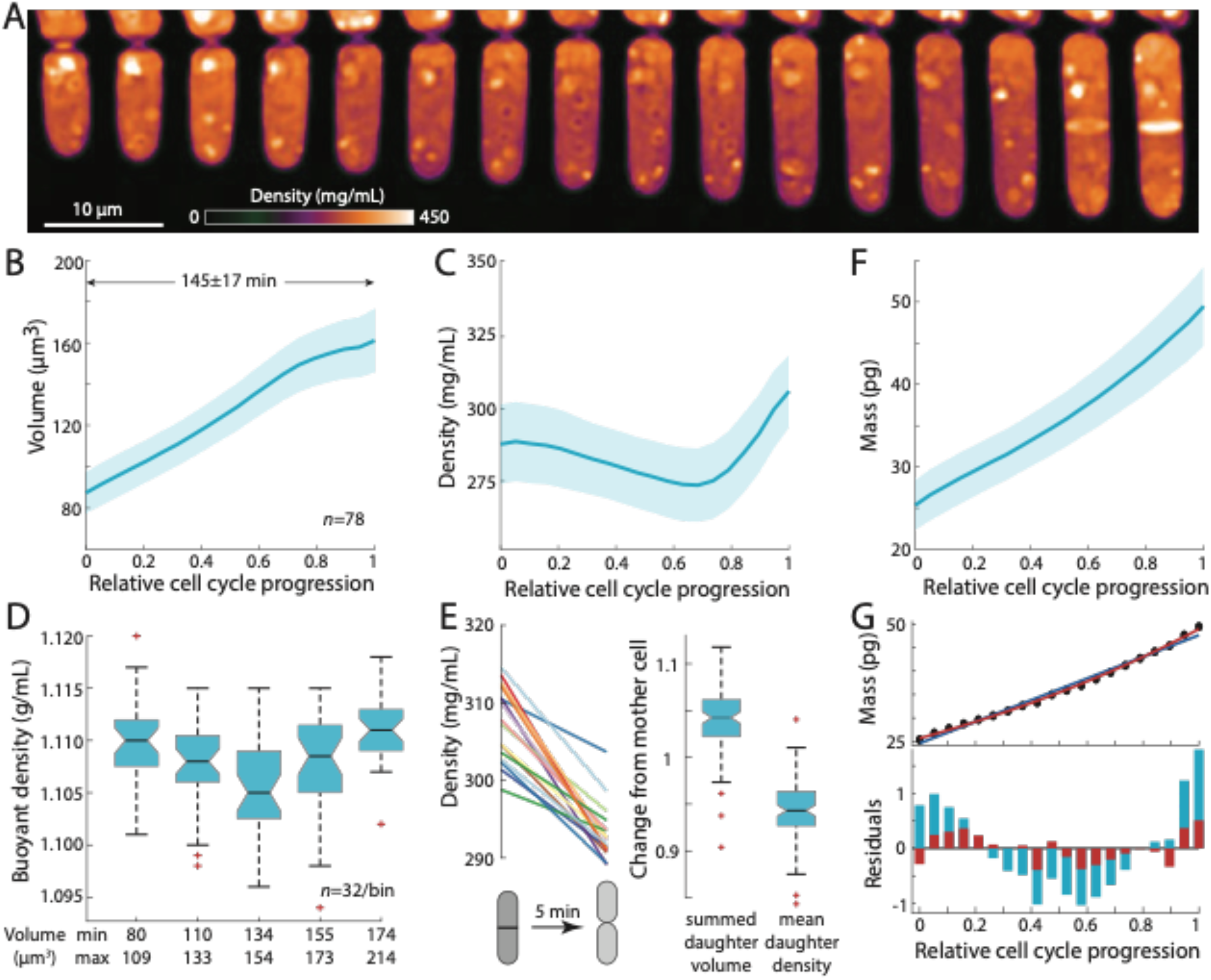
Intracellular density varies across the cell cycle. A) Wild-type fission yeast cells in exponential phase were imaged in time lapse in a microfluidic chamber and phase-shift maps were extracted by QPI. Shown are images of a representative cell traversing the cell cycle from (left) cell birth to (right) septation (10 min/frame). B,C,F) Cell volume (B), density (C), and dry mass (F) of cells aligned by their relative progression in the cell cycle. Curves are mean values and shaded regions represent 1 standard deviation (*n*=78 cells). Mass was estimated from volume and density measurements. D) SMR-based measurements of fission yeast cells in an asynchronous culture (binned by cell volume) showed a similar decrease in buoyant density at intermediate volume as QPI measurements of dry-mass density (C). E) Cell density decreases upon cell separation. Left: density measurements of cells just before and just after cell separation (5 min apart). Right: normalized changes in volume and density between the mother and resultant daughter cells. G) Dry mass grows more exponentially than linearly. The residuals (bottom) of an exponential fit (red) to mass growth were much smaller than a linear fit (blue). See also Figure S2A.

Intracellular density displayed consistent dynamics during the *S. pombe* cell cycle. Density gradually decreased from the beginning of the cell cycle through G2 phase by ~5%, followed by a steady rise during mitosis and cytokinesis (Figure 2C). We confirmed these findings using a complementary approach for measuring density. Holographic imaging (Methods) showed that the mean refractive index of cells decreased with cell length prior to septation, whereas cells of similar length with a septum exhibited higher refractive indices (Figure S2A,B) (17). We used a suspended microchannel resonator (SMR) to measure the buoyant mass of single cells in two different density solutions and thereby derive single-cell buoyant densities and volumes (18) (Methods). SMR measurements revealed that fission yeast exhibit a mean buoyant density of 1.108±0.005 g/mL, and buoyant densities were lower in intermediate-sized cells than in small and large cells (Figure 2D, S2C,D), consistent with our QPI-based observations of cell cycle-dependent densities (Figure 2C, S2E). As expected, the relative changes in buoyant density were much less than changes in dry-mass density due to the different nature of the respective measurements: dry-mass density accounts only for a fraction of buoyant density as cells are comprised mostly of water mostly, and hence relative changes are magnified (19).

We also identified an additional change in density at cell birth. At the end of cytokinesis, the middle layer of the septum is digested, and daughter cells separate (20, 21). During this 5-10-min period, the septum bulges outward on each side to form the rounded shape of the new end, in a physical process driven by turgor pressure (22). QPI analysis of individual cells showed a consistent drop in density within a 5-min window around cell separation (Figure 2E, S3); during this period, cell volume increased by ~5% while density decreased by ~5% (Figure 2E). The magnitude and rapidity of the density change suggest that the volume increase at cell separation is due primarily to swelling from water uptake. Thus, density appears to be directly tied to the dynamics of volume expansion.

Our precision measurements of cellular dimensions and intracellular density provide a quantitative characterization of dry-mass dynamics (biosynthesis) throughout the cell cycle. The absolute rate of dry-mass accumulation steadily increased during the cell cycle (Figure 2F). Dry-mass dynamics were more exponential than linear in nature (Figure 2G, S4A), as shown by a comparison of linear versus exponential fits (Figure 2G) and in the dynamics of normalized mass growth (Figure S4A). The absolute rate of mass synthesis was therefore higher during mitosis and cytokinesis than at the beginning of the cell cycle, even though the cell slowed in volume growth late in the cycle. Thus, mass production was not tightly coupled to volume growth. These measurements suggest a simple model in which the increase of density in mitosis and cytokinesis arises as consequence of continued mass accumulation when volume growth is halted.

### Cell-cycle perturbations exacerbate cell cycle-dependent density variation

Our data demonstrate that intracellular density increases during mitosis and cytokinesis as a result of biosynthesis continuing unabated while volume growth slows; conversely, density decreases during interphase because the rate of volume growth surpasses the rate of mass synthesis. However, the origin of these dynamics remains unresolved. One possibility is that the relative rates of mass synthesis and volume growth are not directly coupled; with mass growing exponentially, density variations then arise indirectly as a consequence of cell-cycle regulation of volume expansion, which is controlled by cell polarity programs that redirect the cell wall growth machinery to the middle of the cell for septum formation prior to cytokinesis (23–25). Alternatively, the density at each cell-cycle stage could be directly programmed to specific levels by specific cell-cycle regulators. Another possibility is that density variations may be due to a cell cycle-independent oscillator, such as a metabolic oscillator (26). To distinguish these models, we examined the consequences of arresting or delaying cells at particular stages of the cell cycle (Figure 3A). If there is no strict control of biosynthesis, then when mitosis or cytokinesis is blocked, mass should continue to accumulate and density should reach higher levels than in normal cells, and conversely density should fall below normal in extended interphase. If density is instead regulated at specific levels according to cell-cycle phase, density levels should not change during cell-cycle delays beyond the ranges appropriate for each phase. If a cell cycle-independent oscillator governs density variations, then oscillations could continue even during cell-cycle arrests (27, 28).

**Figure 3:**
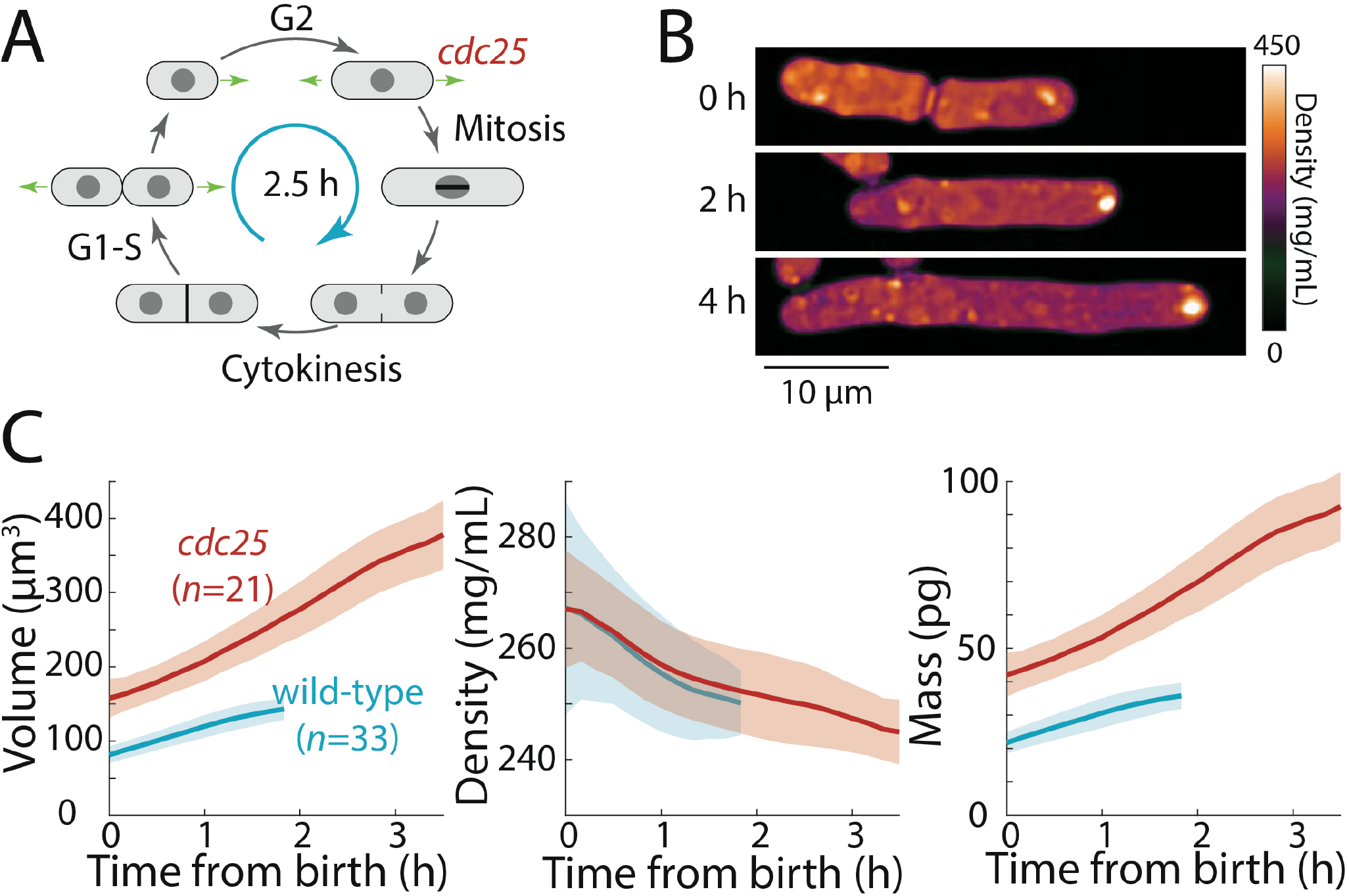
Extension of the G2 phase of the *S. pombe* cell cycle results in cell elongation and decreased intracellular density. A) Schematic of the fission yeast cell cycle, highlighting the point at which the *cdc25* temperature-sensitive mutant delays the G2-M transition. B) *cdc25-22* cells were shifted from the permissive temperature 25 °C to the semi-permissive temperature 32 °C to extend G2 phase, leading to continued cell elongation. QPI density maps of a representative cell are shown. Note that density decreased during cell elongation. C) QPI-based measurements of volume (left), density (middle), and dry mass (right) of *cdc25-22* cells (red) that grew at least 2.5-fold relative to their birth length before dividing, compared with wild-type cells (blue) under the same conditions. Measurements are aligned from cell birth until elongation rate decreased to 20 nm/min (as an indication of the transition to mitosis). Curves are mean values and shaded regions represent 1 standard deviation.

First, we tested whether density would decrease further in *S. pombe* cells experiencing an extended period of growth during interphase. We delayed cells harboring a *cdc25*-*22* mutation (29) in G2 phase by shifting them from room temperature to the semi-permissive temperature (32 °C). These cells continued to grow from their tips and formed abnormally elongated cells (Figure 3B,C) before dividing. To focus on cells that remained in G2 phase for an extended interval, we limited our analysis to cells that elongated to >2.5-fold their initial length. In these mutant cells, during their prolonged G2 phase of 2-3 h, intracellular density decreased further than in wild-type cells (~8% in *cdc25-22* cells from 267±19 to 245±6 mg/mL, compared to ~5% in wild-type cells from 267±11 to 250±6 mg/mL) (Figure 3C). No evidence of density oscillations in the prolonged G2 phase was evident, arguing against cell-cycle independent oscillations as the cause of density variations during a normal cell cycle. These data suggest that density falls during G2 phase because the rate of volume growth continues to be slightly faster than the rate of biosynthesis.

Second, we tested whether cells arrested in mitosis displayed increased intracellular density (Figure 4A). We delayed cells in metaphase using a *cut7-ts* mutant (kinesin-5) defective in mitotic spindle assembly (30). Using QPI, we tracked intracellular density from mitosis initiation until the earliest signs of septum formation. As expected, at the non-permissive temperature this interval was longer for *cut7-ts* cells (20-30 min) than for wild-type cells (10-20 min; Figure 4B,C). During this mitotic period, the density of wild-type and *cut7-ts* cells increased at a similar rate (Figure 4B,C): hence, the extended time in metaphase in *cut7-ts* cells led to a greater density increase (7% in *cut7-ts* from 270±12 to 288±14 mg/mL versus 5% in wild-type cells from 264±11 to 278±13 mg/mL).

**Figure 4:**
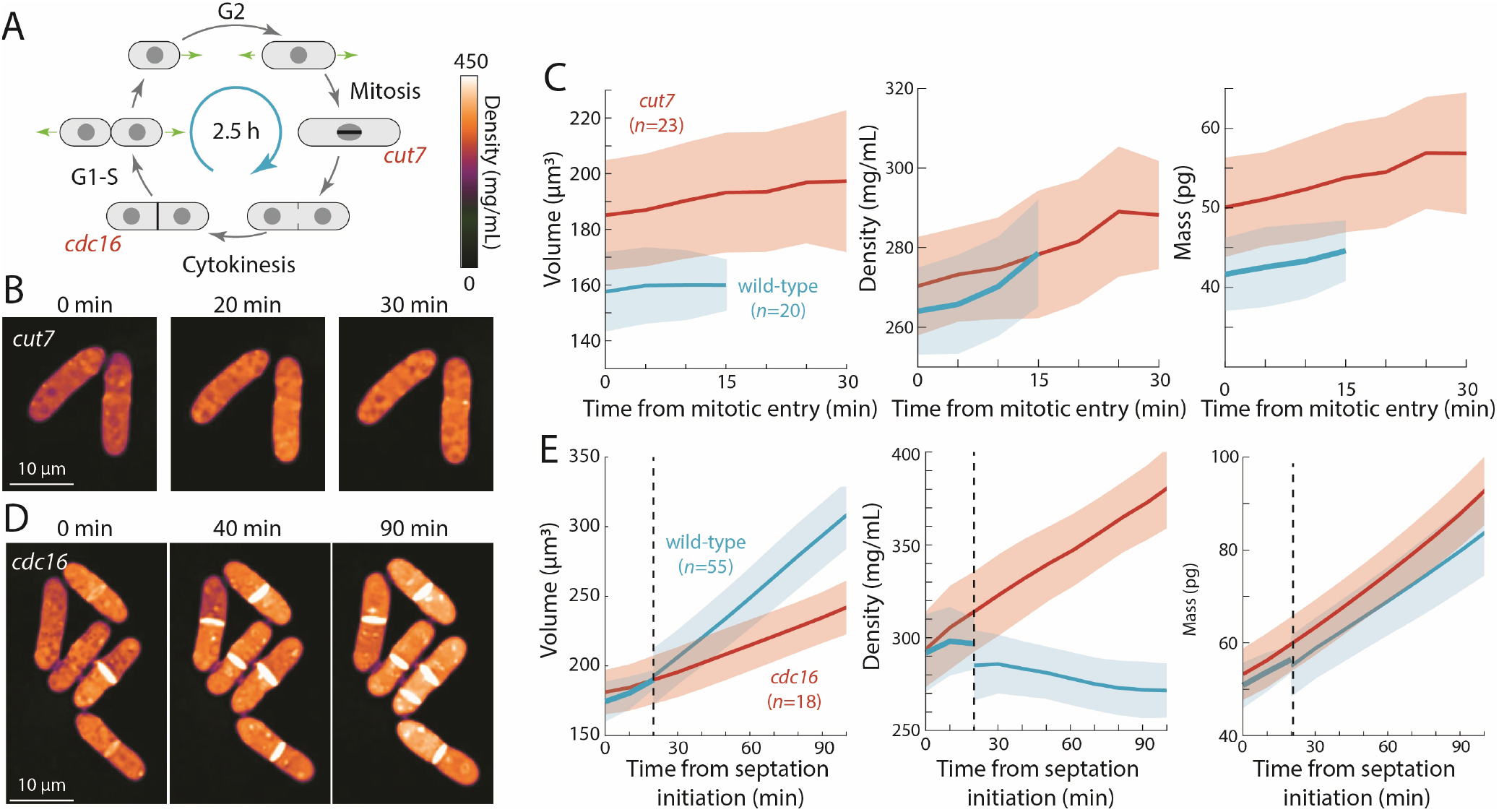
Cell-cycle arrests in mitosis and cytokinesis result in increased intracellular density in *S. pombe*. A) Temperature-sensitive mutants *cut7-446* (spindle kinesin-5) and *cdc16-116* block the cell cycle in mitosis and cytokinesis, respectively. B) *cut7-446* cells were shifted from 25 °C to 30 °C to delay mitotic progression. QPI of two representative cells delayed in mitosis for ~20 min until the onset of septation (30 min time point). Note that density continued to increase during mitotic arrest. C) QPI-based measurements of volume (left), density (middle), and dry mass (right) of *cut7-446* cells from mitotic entry (*t*=0) through initiation of septum formation at cytokinesis. Curves are mean values and shaded regions represent 1 standard deviation. D) *cdc16-116* cells were shifted from 25 °C to 34 °C to arrest cells in cytokinesis. QPI of five representative cells are shown. *cdc16* cells generally did not complete cell separation and often assembled additional septa without elongating. Note that density increased during this cytokinetic arrest. E) QPI-based measurements of volume (left), density (middle), and dry mass (right) of *cdc16-116* cells from initiation of the first septum (*t*=0). Wild-type cells separated after ~20 min (dashed line), and thereafter the behavior of the daughter cells was tracked for comparison with *cdc16* cells (volume and dry mass were summed for the two daughter cells). Curves are mean values and shaded regions represent 1 standard deviation.

Third, we arrested cells in cytokinesis, again to test for an increase in density (Figure 4A). *cdc16-116* mutant cells arrest in cytokinesis, and thus repeatedly make septa without elongating (31). Upon a shift from 25 °C to the non-permissive temperature (34 °C), cells that maintained cytokinetic arrest continued to increase in cytoplasmic density; density after 90 min was 20-30% higher than in cytokinesis-competent wild-type cells (Figure 4D,E). Thus, biosynthesis continues throughout an extended block of mitosis or cytokinesis, leading to abnormally high intracellular density.

Finally, we asked whether inhibition of volume growth is sufficient to increase cytoplasmic density. We previously showed that two treatments that slow volume growth (osmotic oscillations and treatment with brefeldin A) led to an increase in cytoplasmic density (10). However, since these treatments do not completely halt volume growth and/or result in cell death, we treated wild-type cells with the F-actin inhibitor latrunculin A, which causes immediate cessation of tip growth independent of cell-cycle stage (32, 33). QPI density maps showed that latrunculin A treatment caused all cells to immediately halt tip growth and begin to steadily increase in density, regardless of cell-cycle stage (Figure 5A,B). The mean density increase after 1 h was ~20% (Figure 5B). Similar increases in density were seen in cells of different sizes (Figure S5). However, we noted that in contrast to the mitotic and cytokinesis arrests, mass increases were variable and on average increased more slowly during latrunculin A treatment than during normal growth (Figure 2F), suggesting a partial slowdown in biosynthesis and/or an increase in degradation.

**Figure 5:**
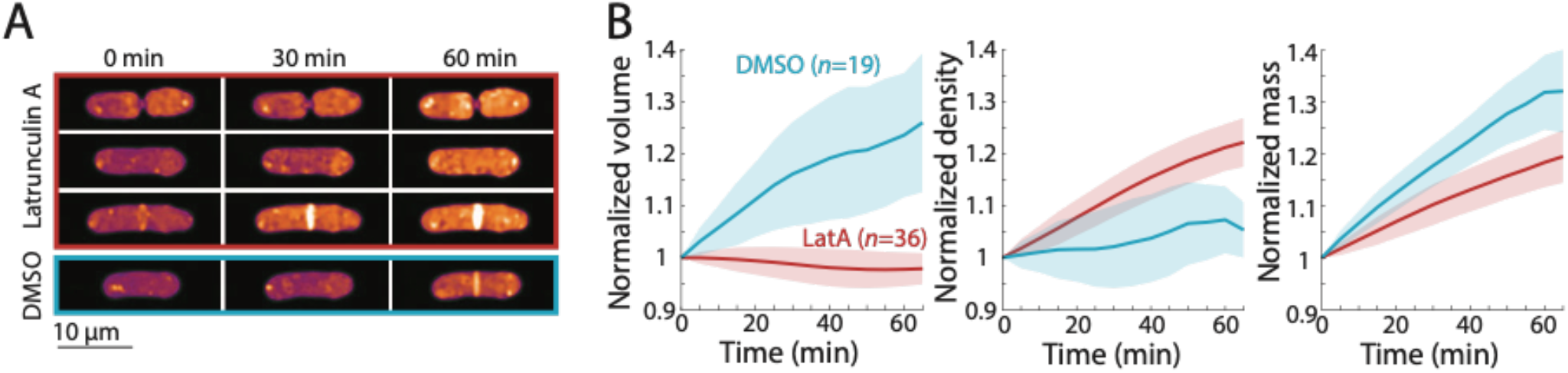
Cell cycle-independent growth inhibition by latrunculin A results in increased intracellular density in *S. pombe*. A) Latrunculin A treatment inhibited cell growth and increased intracellular density regardless of cell-cycle stage. Representative QPI density maps of three wild-type cells at different points of the cell cycle treated with 200 μM latrunculin A for the indicated times. As a control, cells were treated with 1% DMSO; growth continued and density remained relatively constant. B) QPI-based volume (left), density (middle), and dry mass (right) measurements of latrunculin A (LatA)-treated or DMSO-treated wild-type cells from the start of treatment (*t*=0). Growth halted and density increased due to continued mass synthesis during treatment. Curves are mean values and shaded regions represent 1 standard deviation.

Taken together, these results show that cell density differences are exacerbated during cell cycle arrests and that inhibition of volume growth is sufficient to increase intracellular density. These findings are not consistent with models in which density is set at cell-cycle stage-specific levels or involving cell cycle-independent oscillators. Rather, our results strongly support a model in which the variations in intracellular density arise from cell cycle-dependent changes in volume growth rate.

### A polarized density gradient is associated with the pattern of tip growth

Fission yeast have a well-known pattern of volume growth in which after cell division, the old end initiates tip growth soon after cell birth, and partway through G2, tip growth at the new end begins, but at a slower rate than the old end (16, 34). As expected, our time-lapse data showed that the old and new ends grew over the course of the cycle on average by ~4 and 2 μm, respectively. In QPI density maps, we noted that many cells exhibited a gradient of intracellular density in which the ends that were actively growing appeared less dense than the non-growing ends (Figure 6A). We hypothesized that these subcellular gradients reflected differences in tip growth between the two ends of the cell. In agreement with our hypothesis, the slower-growing new end typically appeared denser than the faster-growing end (Figure 6A). In some cells, the difference in densities between the fast- and slow-growing ends was ~10% of the mean overall density (Figure 6A). The mean density difference between the two ends throughout the cell cycle was ~15 mg/mL, corresponding to ~5% of the mean overall density (Figure 6B). To address the potential for differences in the widths (and hence heights above the coverslip) of old and new ends to influence phase shifts (Figure S6A), we constrained our analysis to cells within a narrow range of widths and found that local density and tip growth remained highly correlated (Figure S6B). Examples of post-cytokinesis cells with adjacent compartments exhibiting different densities (see Figures 7, S7 below) further indicated that these differences could not be explained simply by width differences. Since tip growth is regulated by actin-dependent mechanisms (32–34), we tested whether maintenance of the gradient is dependent on F-actin or tip growth: we found that in cells treated with latrunculin A, spatial density gradients persisted over time (Figure S6C). These results demonstrate that intracellular density gradients are stable and linked to local growth patterns, and that their maintenance does not require active growth or F-actin.

**Figure 6:**
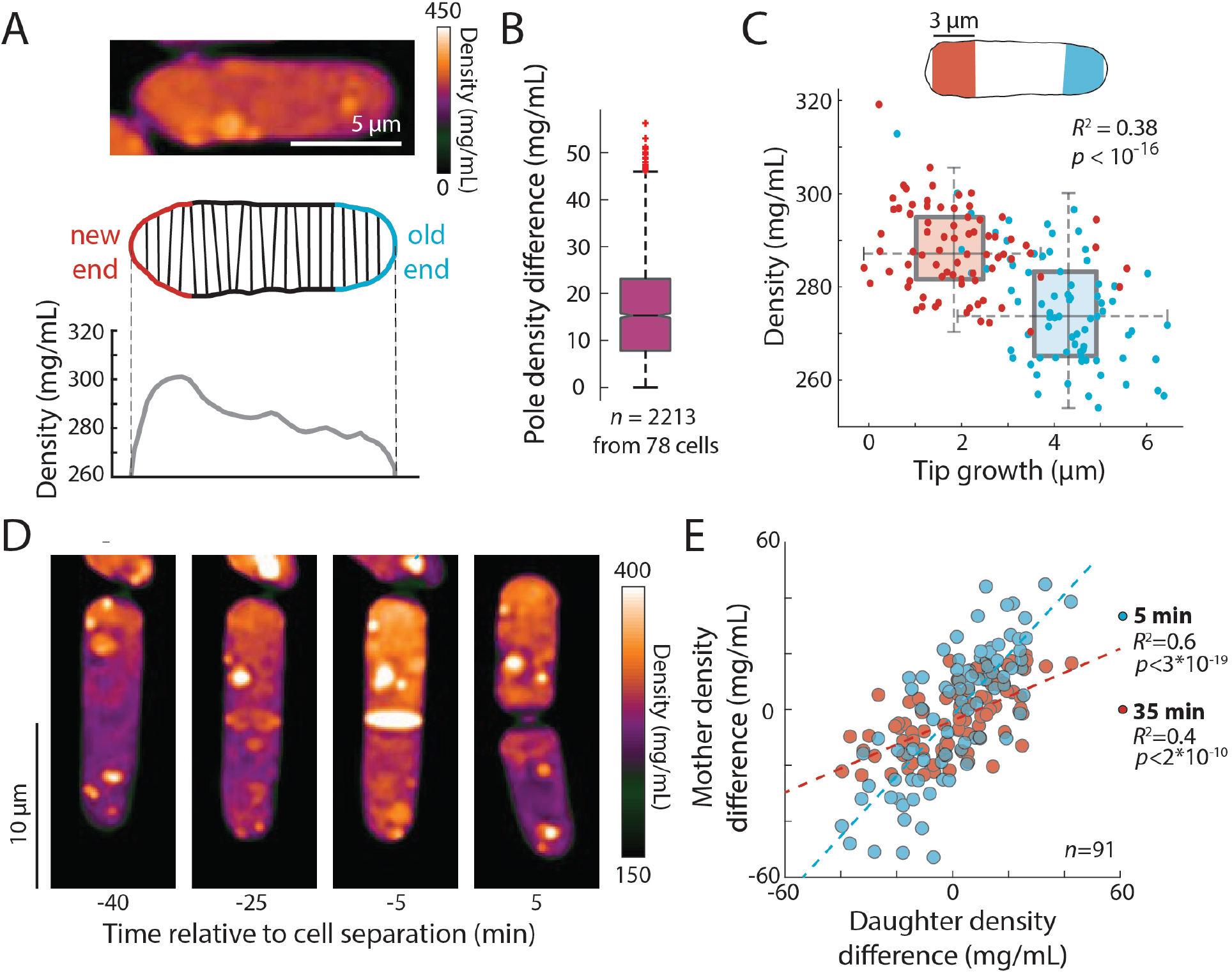
An intracellular density gradient negatively correlates with tip growth in *S. pombe*. A) Top: QPI of a representative cell displaying an intracellular gradient of density. Middle: Density was measured in slices perpendicular to the long axis. Bottom: The new end (non- or slowly growing) exhibited a higher density than the old (growing) end. B) Density was substantially different between the new and old ends in many cells. Time-lapse QPI was used to measure the densities in regions within 3 μm of each cell end. Shown is the density difference between the cell poles averaged over the cell cycle. Box extends from 25^th^ to 75^th^ percentile, with the median as a horizontal bar. Whiskers indicate extreme points not considered outliers (*n*=78 cells). C) Mean density and amount of tip growth over an entire cell cycle was measured in 3 μm regions at the old (blue) and new (red) ends. Old ends grew more and exhibited lower mean densities over the course of the cell cycle than new ends. D) QPI density map of a representative cell at interphase, the start of septum formation, late in septum formation, and after cell division. The gradient of intracellular density in the interphase cell was maintained over time and passed on to the daughter cells. E) Asymmetric density patterns are propagated to the next cell cycle. The density difference between the two ends of a mother cell correlated with the density difference between the progeny daughter cells. Shown are Pearson’s correlation coefficient between the density difference of daughter cells and the corresponding halves of the mother cell at 5 min (blue) or 35 min (red) before cell division. The halves of the mother cell exhibited a larger range of density differences at the later time point, when they were more consistent with the density differences between daughter cells.

**Figure 7:**
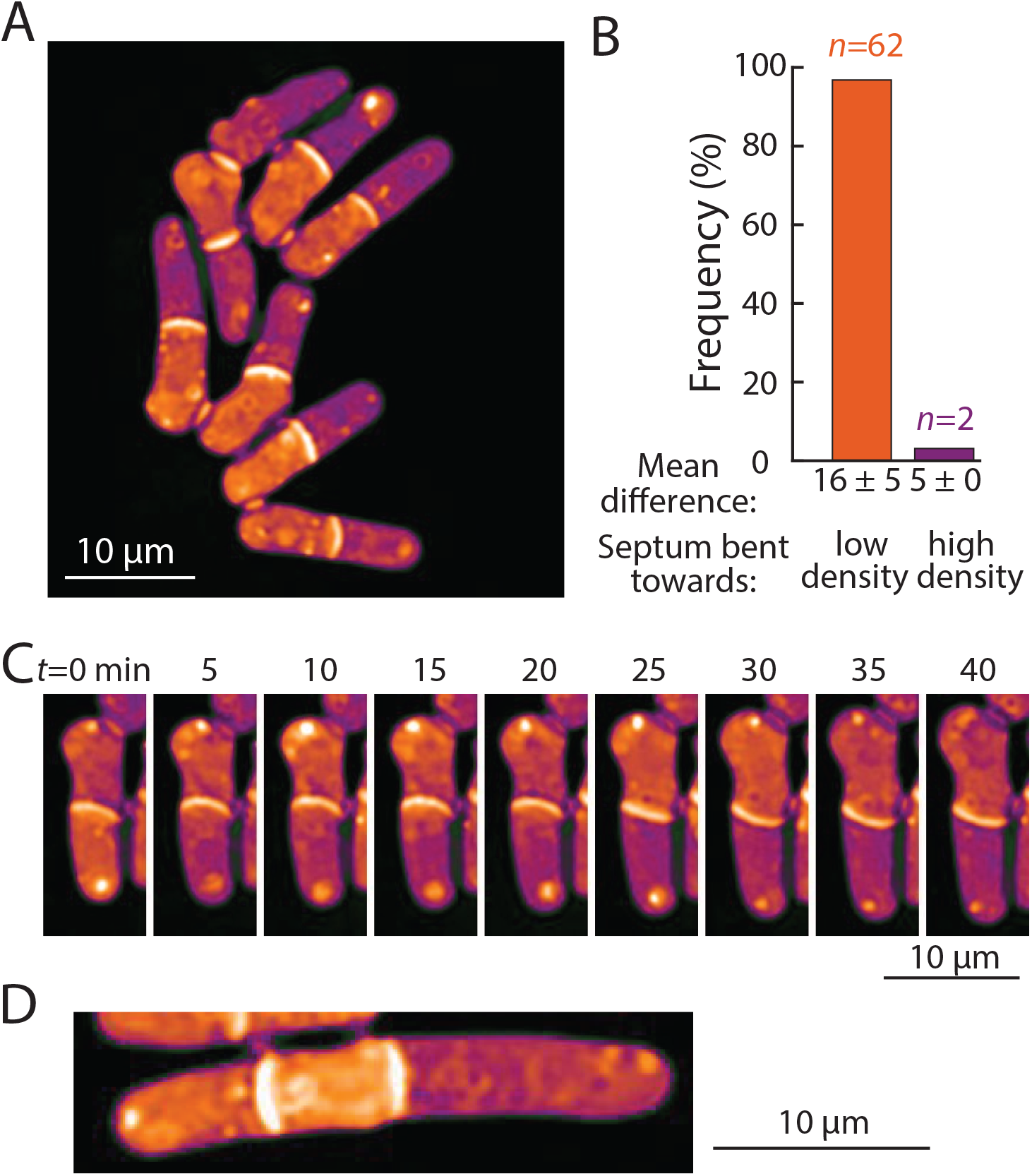
Bending of the septum reveals a link between intracellular density and osmotic pressure. A) QPI density map of *mid2Δ cells* that are delayed in cell separation showed bent septa and density differences between two daughter-cell compartments. Note that the septa are bent away from the denser compartment in all cells in this image. B) The direction of septum bending was almost always toward the lower-density daughter cell. The direction of the bent septum was measured at the time of maximum density difference between daughter-cell compartments. C) An example of a cell in which the direction of septal bending and the sign of the density difference between daughter-cell compartments fluctuated over time. After the bottom compartment decreased and the top compartment increased in density, the septum bent in the opposite direction, consistent with the correlation between bending and density difference in (B). D) In multi-septated *mid2Δ* cells, internal compartments bounded by two septa exhibited higher density than the surrounding compartments; in these situations, both septa typically bent away from the higher-density compartment.

We noticed asymmetries in density between daughter cells after cell division, in which one of the daughters was denser than its sister. Time-lapse imaging showed that the intracellular density differences established during interphase were often propagated through cell division and correlated with density differences between the progeny daughter cells after cytokinesis (Figure 6D,E). Thus, subcellular density variations are stable enough to be propagated through generations.

### Cell density differences are linked to intracellular osmotic pressure

Next, we ascertained whether density variations of 5-20% – the magnitude observed within and across cells with normal physiology – have physiological consequences. One variable potentially connected with intracellular density is macromolecular crowding. High concentrations of macromolecules are predicted to produce colloid osmotic pressure that may influence cell mechanics (2). As noted above, the densities of daughter compartments in septated cells were often different from each other (Figure 6D,E); these differences were often exacerbated in cells with cell-division defects, such as *mid1*, *mid2*, and *cdc16* cells. We noted that the septum between these daughter cells generally bent away from the more dense compartment (Figure S7). Previous studies (22) reported that the septum is an elastic structure that can be used as a biosensor that reports on osmotic pressure differences between compartments. For instance, when one daughter is lysed by laser microsurgery and loses turgor pressure, the septum bulges away from the intact daughter (22). Temporal fluctuations in septum bending have been proposed to arise from fluctuations in pressure differences between daughter cells (35).

To investigate the relationship between density differences and septum bending, we focused on *mid2Δ* cells. Mid2 is an anillin ortholog that regulates septins in late cytokinesis; *mid2Δ* mutants exhibit long delays (1-2 h) in cell separation and thus most cells in the population have one or more septa (36). We used QPI to track intracellular density over time and identified the time after septum formation at which the maximum density difference was reached for each cell. Of the 71% (64/90) of cells exhibiting a bent septum, 97% (62/64) exhibited a septum bent away from the compartment of higher density at the time of maximum density difference (Figure 7A), with a mean maximum difference of 16±5% (Figure 7B). In 2/90 cells, the septum was bent in the opposite manner, toward the compartment of lower density; in these rare instances, the maximum density difference was substantially lower (5%; Figure 7B) and may represent cases in which the septum is fluctuating in direction. Indeed, in one cell the fluctuating direction of septum bending coincided with alternation of the sign of the density difference between the daughter cells (Figure 7C). In instances where the septum appeared flat, the density difference was substantially lower (~4.5%) than in cells with a bent septum.

We also noted density variations and bent septa in multi-septated *mid2Δ* cells. In particular, internal compartments bounded by two septa were hampered in their ability to grow in volume and correspondingly exhibited higher density than the surrounding compartments (Figure 7D) These situations were frequently associated with both septa bending away from the higher-density compartment (Figure 7D). The observation that septal bending occurred for density differences as low as 5-10% suggests that the density variations over the course of a normal cell cycle (Figure 2), in cell cycle arrested cells (Figure 3, 4, 5) or between growing and non-growing cell tips (Figure 6) may reflect substantial changes in turgor-mediated stresses.

## Discussion

Here, we established a QPI method based on *z*-stacks of brightfield images for quantifying intracellular density dynamics in living cells. We found that exponential-phase fission yeast cells have a dry-mass density of 282±16 mg/mL (Figure 1C-E) and a buoyant-mass density of 1.108±0.005 g/mL (Figure 2D), comparable to measurements in other organisms (3, 37). Density varied systematically across the cell cycle in wild-type fission yeast cells over a range of ~10% (Figure 2C), while the relative rate of dry-mass synthesis remained constant (reflecting exponential accumulation) throughout all cell-cycle stages (Figure 2F,G). These quantitative findings, which utilize precise sub-pixel measurements of cellular dimensions and automated analysis platforms, are consistent with more qualitative density studies of fission yeast using other methods (17, 38, 39).

Our data support a model in which these density variations during the cell cycle are a product of programmed changes in volume growth accompanied by a constant relative rate of mass biosynthesis. Accordingly, volume growth and biosynthesis were not tightly coupled throughout the cell cycle. During tip growth in G2 phase, intracellular density dropped steadily (Figure 2C, 3C), indicating that the rate of volume growth outpaces biosynthesis during this period. Density steadily rose during mitosis and cytokinesis (Figure 2C), when volume growth ceases or slows. After separation of daughter cells (cell birth), density dropped during the rapid increase in cell volume as the new cell end expanded (Figure 2E), possibly due to water influx. Despite these changes in growth rate and density, it is remarkable that even without tight feedback controls, cells maintained a relatively tight distribution (CV~6%) of densities across the population (Figure 1C).

Consistent with this model, density shifts were exacerbated by perturbations of cell-cycle progression or of volume growth directly. *cdc25* mutants delayed in G2 phase exhibited a steady decline in density as cells elongated abnormally (Figure 3), reminiscent of the cytoplasmic dilution observed in very enlarged budding yeast cells and senescent mammalian cells (9). During mitotic arrest at the spindle checkpoint (*cut7*) or cytokinesis arrest through regulation of the SIN pathway (*cdc16*), volume growth was slowed but mass synthesis was unaffected, resulting in steady density increases (Figure 4C,E). Inhibition of cell growth with latrunculin A caused a steady increase in density regardless of cell cycle stage (Figure 5). These findings suggest that any perturbation that affects cell-cycle progression or growth is likely to alter density dynamics. Cell-cycle arrests are commonly used to synchronize cells and are often triggered in response to stresses such as DNA damage. Our study demonstrates that such perturbations are not as innocuous as often thought; arrests not only affect cell-cycle progression and cell size, but also cause changes in intracellular density and potentially other downstream changes in physiology.

Our studies provide quantitative measurements of the mass dynamics of individual fission yeast cells throughout the cell cycle. We found that mass accumulation was continuous, consistent with previous studies (38, 39). Notably, our quantitative measurements suggested that mass dynamics were exponential in nature (Figure 2F,G, S2F), consistent with findings in other cell types (40). Intriguingly, studies of density and growth in other cell types reported somewhat different cell-cycle patterns. In budding yeast, buoyant density is lowest in early G1 and rises in late G1 and S phase at the time of bud formation (37). Moreover, cell-cycle arrests in S and M phase and latrunculin A treatment do not lead to increases in buoyant density in budding yeast, unlike our findings in fission yeast (Figure 5). In human cells, mass growth continues in early mitosis, but stops in metaphase and resumes in late cytokinesis, potentially with subtle oscillations (41, 42). Density is constant in human cells during much of the cell cycle, but decreases in mitosis (by 0.5% in buoyant mass, equivalent to >10% decrease in dry-mass density) coincident with a 10-30% volume increase during mitotic rounding; density then slightly increases in cytokinesis (6, 7). In the bacterium *Escherichia coli*, density varies somewhat from birth to division, but the ratio of surface area to mass is relatively constant, suggesting that biosynthesis is linked to surface-area synthesis (43). Here we found that in fission yeast, mass was also more closely coupled with surface area than volume, especially when the surface area of both sides of the septum were factored in (Figure S4B-D). However, examples such as latruculin A-treated cells (Figure 5B) demonstrated that mass accumulation and surface area growth are not inextricably linked and can be uncoupled. It remains to be seen whether general rules of cell-density regulation will emerge from comparisons across organisms and cell types.

Our findings also provide insight into the relationship between the rate of volume growth and subcellular density regulation. Fission yeast cells grow through tip growth, which involves the extension and assembly of new cell wall and plasma membrane at the growing tip; this growth is accomplished by a complex integration of the cell-polarity machinery, exocytosis, wall growth and mechanics, and turgor pressure (25, 34). In addition to the global effect of volume growth on density, the intriguing polarization of density patterns that we discovered (Figure 6) suggests that tip growth influences local intracellular density patterns more directly. Spatial gradients revealed that local density was correlated with tip growth, with growing ends having lower density (Figure 6C). It is not yet clear what cellular components are responsible for this spatial pattern, and whether they are actively depleted at growing cell tips or concentrated at non-growing regions. The relevant components may be membrane-bound or membrane-less organelles; it is unlikely that they are soluble, freely diffusing particles, unless a diffusion barrier (perhaps the nucleus) exists. The distributions of large organelles and total protein are not polarized in this manner (44, 45). However, polarized patterns have been detected in immunofluorescence of damaged (carbonylated) proteins, which may impact replicative aging (46). Polarized density patterns established in interphase were often propagated through cell division and appeared to be inherited by the daughter cells, resulting in differences in density across a lineage (Figure 6D,E). How these patterns may lead to asymmetric behaviors in cell lineages such as growth patterns and aging remains to be explored (34, 46). Spatial heterogeneities in density have also been observed in mammalian cells; it remains to be determined whether these patterns are related to cell shape, organelles, or variations in cytoplasmic density (5, 47, 48).

The effects of changes in intracellular density on cellular functions are only beginning to be appreciated. The density changes of 5-20% that we observed, which likely affect the concentration of most if not all cellular components in the cytoplasm, could have profound consequences for the biochemistry of cellular reactions and on macromolecular crowding. Here we provided evidence that density may also affect cell mechanics and thereby influence cell shape. Intracellular osmotic pressure and density differences between daughter compartments were strongly coupled, as evidenced by bending of the elastic septal cell wall (Figure 7). Density may affect pressure through effects on macromolecular crowding, which produces colloid osmotic pressure associated with the displacement of water (2). Differences in colloid osmotic pressure have been proposed to influence nuclear size (2, 49), but experimental evidence linking crowding to force generation *in vivo* generally remains scant. Our findings thus lend important support to the idea that different densities of macromolecules can generate large enough differences in colloid osmotic pressure to alter cell shape. We speculate that the increase of density at cell division may provide mechanical force through increased turgor pressure to facilitate cell-cell separation and bulging of the cell wall (22), and may affect the distribution of mechanical stresses within the cell wall during other cell-cycle stages as well.

Another possible consequence of intracellular density changes is the regulation of volume growth rate. In fission yeast, perturbations that increase density are followed by a dramatic increase in volume growth rate that persists for hours (10). How density impacts volume growth rate, possibly via the concentration of certain intracellular factors or through effects on intracellular pressure, remains to be elucidated. Thus, toggling the rate of volume growth may play an important role in density homeostasis. Future studies focusing on the effects of cell density on particular cellular processes will be needed to understand the full scope of consequences of density variations on cell physiology.

## Methods

### Strains and cell culturing

All strains used in this study are listed in Table S1. Methods for propagation and growth of *S. pombe* cells were as described in (50). In general, cultures were grown in 3 mL of YE5S medium at 30 °C on a rotating shaker overnight to OD_600_~1, diluted to OD_600_~0.1, and incubated until OD_600_~0.3 for imaging. Temperature sensitive mutant c*dc25-22* cells (and wild-type control cells) were first grown at room temperature; 90 min after imaging started, the temperature was increased to 32 °C. Temperature-sensitive mutant *cdc16-116* cells (and wild-type control cells) were first grown at 25 °C, then imaged on the microscope with the temperature-controlled enclosure pre-heated to 34 °C. Temperature-sensitive mutant *cut7-446* cells (and wild-type control cells) were first grown at 25 °C, then imaged on the microscope with the temperature-controlled enclosure pre-heated to 30 °C.

### Single-cell imaging

Images were acquired with a Ti-Eclipse inverted microscope (Nikon) equipped with a 680-nm bandpass filter (D680/30, Chroma Technology) in the illumination path with a 60X (NA: 1.4) DIC oil objective (Nikon). Before imaging, Koehler illumination was configured and the peak illumination intensity at 10-ms exposure time was set to the middle of the dynamic range of the Zyla sCMOS 4.2 camera (Andor Technology). μManager v. 1.41 (51) was used to automate acquisition of *z-*stack brightfield images with a step size of 250 nm from ±3 μm around the focal plane (total of 25 imaging planes) to ensure substantial oversampling that facilitated correcting for potential drift in the *z*-direction over the course of each experiment at 5- or 10-min intervals at multiple (*x,y*) positions.

### Microfluidics

Cellasic microfluidic flow cell plates (Millipore, Y04C) controlled by an ONIX or ONIX2 (Millipore) microfluidic pump system were used for imaging. YE5S medium was loaded into all but one of the fluid reservoirs; the remaining well was loaded with 100 mg/mL bovine serum albumin (BSA) (Sigma Aldrich) in YE5S. Liquid was flowed from all 6 channels for at least 5 min at 5 psi (corresponding to 34.5 kPa), followed by 5 min of flow from YE5S-containing wells to wash out buffer and to fill channels and imaging chambers. The plate was kept in a temperature-controlled enclosure (OkoLab) throughout loading. Cells were then transferred into the appropriate well and loaded into the microfluidic imaging chamber such that a small number of cells were initially trapped, and flow of YE5S was applied. To ensure full exchange of liquid in the chamber during imaging, the flow channel was switched at least 40 s before images were acquired. Every ~2 h, BSA flow was activated during one time point of imaging to calibrate QPI measurements.

### Image analysis to retrieve phase shifts

To reduce post-processing time, each *z*-stack was cropped to a square region containing the cell(s) of interest and a border of at least 40 pixels, and the focal plane was identified. This cropping was accomplished first using FIJI v. 1.53c to identify regions of interest (ROIs) within a thresholded standard deviation *z*-projection image of each brightfield *z*-stack. Using custom Matlab R2019a (Mathworks) scripts, images were cropped to the ROIs and the standard deviation of the pixels in each ROI was computed. The focal plane was defined based on the image in the stack with the lowest standard deviation. Three images above and three images below the focal plane separated by 500 nm were used to quantify cytoplasmic density. Based on these images, the phase information was calculated using a custom Matlab script implementing a previously published algorithm (15). In brief, this method relates the phase information of the cell to brightfield image intensity changes along the *z*-direction. Equidistant, out-of-focus images above and below the focal plane are used to estimate intensity changes at various defocus distances. A phase-shift map is reconstructed in a non-linear, iterative fashion to solve the transport-of-intensity equation.

### Cytoplasmic density quantification

Using Matlab, images were background-corrected by fitting a Gaussian to the highest peak of the histogram (corresponding to the background pixels) of the phase-shift map and shifting every pixel so that the background intensity peak corresponded to zero phase shift. These background-corrected phase-shift maps were converted into binary images using watershedding for cell segmentation; where necessary, binary images were corrected manually to ensure accurate segmentation. Binary images were segmented using Morphometrics (52) to generate subpixel-resolved cell outlines.

Each cell outline was skeletonized using custom Matlab code as follows. First, the closest-fitting rectangle around each cell was used to define the long axis of the cell. Perpendicular to the long axis, sectioning lines at 250-nm intervals and their intersection with the cell contour were computed. The centerline was then updated to run through the midpoint of each sectioning line between the two contour-intersection points. The slope of each sectioning line was updated to be perpendicular to the slope of the centerline around the midpoint. Sectioning lines that crossed a neighboring line were removed. Cell volume and surface area were calculated by summing the volume or area of each section, assuming rotational symmetry. Volume and area of the poles were calculated assuming a regular spherical cap.

To convert the mean intensity of the phase-shift within each cell into absolute concentration (in units of mg/mL), the mean of all cells across all time points was first calculated. Then, the decrease in phase shift induced by a prescribed concentration of BSA (typically 100 mg/mL) was defined as the difference between the mean of the phase shifts before and after the BSA imaging time point and the phase shift during the BSA time point. This difference in intensity established the calibration scaling between phase shift intensity and the concentration of BSA (Figure 1B). The cytoplasmic density of each cell was then calculated by dividing the mean phase shift of the cell by the aforementioned scaling factor. The mass of each cell was inferred from its mean density and volume.

### BSA calibration

Channel slides (μ-Slide VI 0.4, ibidi) were treated with lectin (Sigma-Aldrich, L1395) (0.1 mg/mL in water) for ~5 min, washed with YE5S, and cells were added and incubated for ~5 min to allow for attachment. Unattached cells were removed by washing with YE5S. Attached cells were first imaged in YE5S medium. Freshly prepared BSA (Sigma-Aldrich, A3608) was then added to a final concentration of 200 mg/mL. YE5S was added to dilute BSA to the desired concentrations (150, 100, and 50 mg/mL), followed by washout of the BSA and imaging in YE5S.

### Lipid droplet staining

Lipid droplets were stained with the dye BODIPY 493/503 (Thermo Fisher, D3922) (53). Aliquots (10 μL) of 100 mM BODIPY in absolute ethanol were prepared. Ethanol was then evaporated in a desiccator under vacuum and dried aliquots were stored at 4 °C for long-term storage. For use, an aliquot was redissolved in 10 μL absolute ethanol and 1 μL was added to 1 mL of cell culture in YE5S for each unit of cell density with OD_600_=0.1 and incubated protected from light for ~1 min at room temperature. Cells were then pelleted at 0.4 rcf in a microfuge for 1 min and medium was exchanged with fresh YE5S. Cells were spotted onto agarose pads and imaged with an EM-CCD camera (Hamamatsu) through a spinning-disk confocal system (Yokogawa CSU-10) attached to one of the ports of a Ti-Eclipse inverted microscope with a 488-nm laser. In parallel, brightfield *z*-stack images were acquired for QPI.

### Lineage tracking for time-lapse imaging datasets

First, each cell present at the beginning of the experiment was linked to the closest cell in the next frame based on the distance between centers and the difference in their size (cross-sectional area). A cell was considered the same if the centers between consecutive time points were within 20 pixels (~2 μm) and the cross-sectional area was >70% of the area at the previous time point. This process was iterated to define the lineage until either requirement was violated (usually due to cell division), at which point a new lineage was initialized using the earliest unassigned cell. All lineages were visually inspected and corrected when necessary.

### Polar growth and density quantification

To separately quantify the growth of the new and old ends, fiduciary markers such as birth scars on the cell outline were identified from which the distance to each pole at the beginning and completion of the cell cycle was measured. The density of each polar region was calculated by extracting the peak of the histogram of density values in the region within 3 μm of the pole at each time point, and then calculating the mean over time points.

### Suspended microchannel resonator (SMR) measurements

SMR-based density and volume measurements were carried out according to a previously reported fluid-switching method (18). Briefly, the SMR measures the buoyant mass of a cell by flowing in culture medium through a vibrating cantilever and measuring changes in vibration frequency. The cell is then mixed with a denser medium composed of 50% culture medium and 50% OptiPrep (Sigma-Aldrich), and flowed back through the cantilever to obtain a second buoyant mass measurement 10 s later. Cell volume and density are calculated from the two consecutive buoyant mass measurements based on the known densities of the two fluids (18). Each cell is serially flushed into the system in culture medium from a reservoir at 30 °C. After every hour of measurement, the reservoir is replenished from an exponential-phase culture.

SMR devices were fabricated at CEA-LETI (Grenoble, France). The physical dimensions and operation of the SMR, as well as data analyses, were identical to those reported in (42, 54). Briefly, the SMR cantilever was vibrated in the second flexural bending mode using a piezo-ceramic plate underneath the SMR chip. The vibration frequency of the cantilever was measured using piezo-resistors at the base of the cantilever. A digital control platform was used to drive the cantilever in a feedback mode, where the vibration frequency signal acquired from the piezo-resistor was delayed, amplified, and used as the drive signal to actuate the cantilever. Fluid flow was controlled using two electronic pressure regulators and solenoid valves, which were used to pressurize vials containing the culture medium. A typical cell transit time through the cantilever was 150 ms. System temperature was controlled by mounting the SMR and culture-medium vials on copper stands connected to a heated water bath. All SMR operations were controlled using National Instruments control cards and custom LabVIEW (2012) code. Frequency data were analyzed using previously reported custom Matlab code that measures the maximum frequency change during the transit of each cell through the cantilever (54). Frequency measurements were calibrated using polystyrene beads and NaCl solutions of known density.

### Holographic refractive index measurements

For refractive index measurements, wild-type *S. pombe* cells grown at 30 °C were immobilized on a lectin-coated glass-bottom 35 mm diameter μ-dish (ibidi). Holographic refractive index measurements were acquired with a 3D Cell Explorer system (Nanolive) with a temperature-controlled enclosure set to 30 °C. First, sum images of *z-*stacks of three-dimensional refractive index maps were generated to retrieve cell outlines by watershedding. Cells oriented at an angle to the flat glass bottom dish were ignored. For each remaining cell, the mean refractive index was extracted from each image in the *z*-stack using Matlab and the highest value (assumed to correspond to the middle plane) was used for further analysis.

### Lipid droplet staining

Lipid droplets were stained with the dye BODIPY 493/503 (Thermo Fisher, D3922) (53). Aliquots (10 μL) of 100 mM BODIPY in absolute ethanol were prepared. Ethanol was then evaporated in a desiccator under vacuum and dried aliquots were stored at 4 °C for long-term storage. For use, an aliquot was redissolved in 10 μL absolute ethanol and 1 μL was added to 1 mL of cell culture in YE5S for each unit of cell density with OD_600_=0.1 and incubated protected from light for ~1 min at room temperature. Cells were then pelleted at 0.4 rcf in a microfuge for 1 min and medium was exchanged with fresh YE5S. Cells were spotted onto agarose pads and imaged with an EM-CCD camera (Hamamatsu) through a spinning-disk confocal system (Yokogawa CSU-10) attached to one of the ports of a Ti-Eclipse inverted microscope with a 488-nm laser. In parallel, brightfield *z*-stack images were acquired for QPI.

### Latrunculin A treatment

Stock solutions were made by dissolving 100 μg latrunculin A (Abacam, ab144290) in dimethyl sulfoxide (DMSO, Sigma-Aldrich) to a concentration of 20 mM and stored at −20 °C in 1-μL aliquots. To prepare agarose pads, 1 μL of 20 mM latrunculin A or 1 μL of DMSO was mixed with 100 μL of YE5S medium containing 2% (w/v) agarose UltraPure agarose (Invitrogen Corporation) kept in a water bath at ~70 °C. The mixture was pipetted onto a microscope glass slide and quickly covered with another slide to form flat agarose pads with thickness of ~2 mm. Once pads had solidified, one slide was carefully removed and 1-2 μL of exponential-phase wild-type cells were deposited on the agarose pad. Cells were allowed to settle for 1-2 min before a coverslip was placed on top and sides were sealed with Valap (1:1:1 vaseline:lanolin:paraffin) to prevent evaporation during imaging.

### Statistical analyses

The magnitude of the correlation between two continuous variables was reported using Pearson’s correlation coefficient. *R*^2^ and associated *p*-values were calculated using the built-in function corrcoef in Matlab R2019a (Mathworks).

## Acknowledgments

We thank the Chang and Huang labs for discussion and support, Gabriella Estevam for her contributions to the project at its early stages, Scott M. Knudsen for assisting with cell culturing for SMR measurements, and Sophie Dumont and her lab for discussion. P.D.O. was supported by postdoctoral fellowships from the Swiss National Science Foundation under Grants P2ELP3_172318 & P400PB_180872. F.C. was supported by NIH GM056836. T.P.M. received funding from the Wellcome Trust (110275/Z/15/Z). K.C.H. is a Chan Zuckerberg Biohub investigator.

## Supplemental Table

**Table S1:** *S. pombe* strains used in this study.

| Genotype | Strain number | Source |
| --- | --- | --- |
| *h*- wild-type strain 972 | FC15 | Chang lab |
| *h*- *cdc25-22* | FC342 | Chang lab |
| *h*- *cut7-446 leu1-32* | FC1455 | Chang lab |
| *h*- *cdc16-116* | FC13 | Chang lab |
| *h*- *mid2::kanMX ade6 leu1-32 ura4-D18* | FC881 | Chang lab |

## Supplemental Figures

**Supplemental Figure 1:**
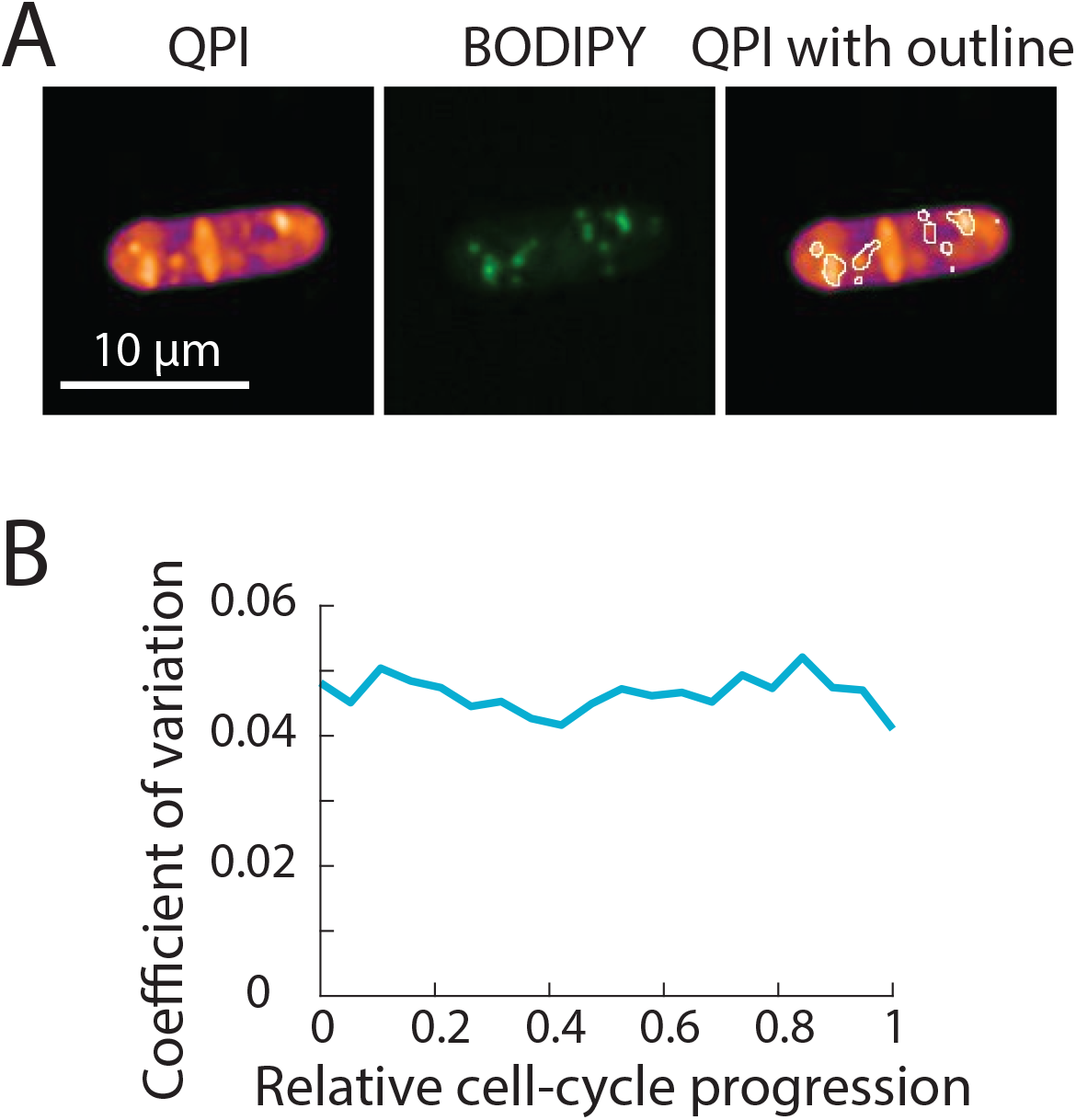
High-density features in QPI co-localize with lipid droplets. A) Cells were stained with BODIPY dye to identify lipid droplets. QPI density map (left) and the corresponding fluorescence image of a representative BODIPY-stained wild-type fission yeast cell (middle). The fluorescence intensity image was thresholded to identify regions containing lipid droplets (outlines), which overlapped with high-density regions in QPI (right). B) Coefficient of variation (CV) of intracellular density over the cell cycle. The CV at each specific stage of cell-cycle progression (<5%) was slightly lower than the CV across an entire cell population (6%, Figure 1C), supporting the notion that some population-wide variation arises from cell cycle-dependent density variations.

**Supplemental Figure 2:**
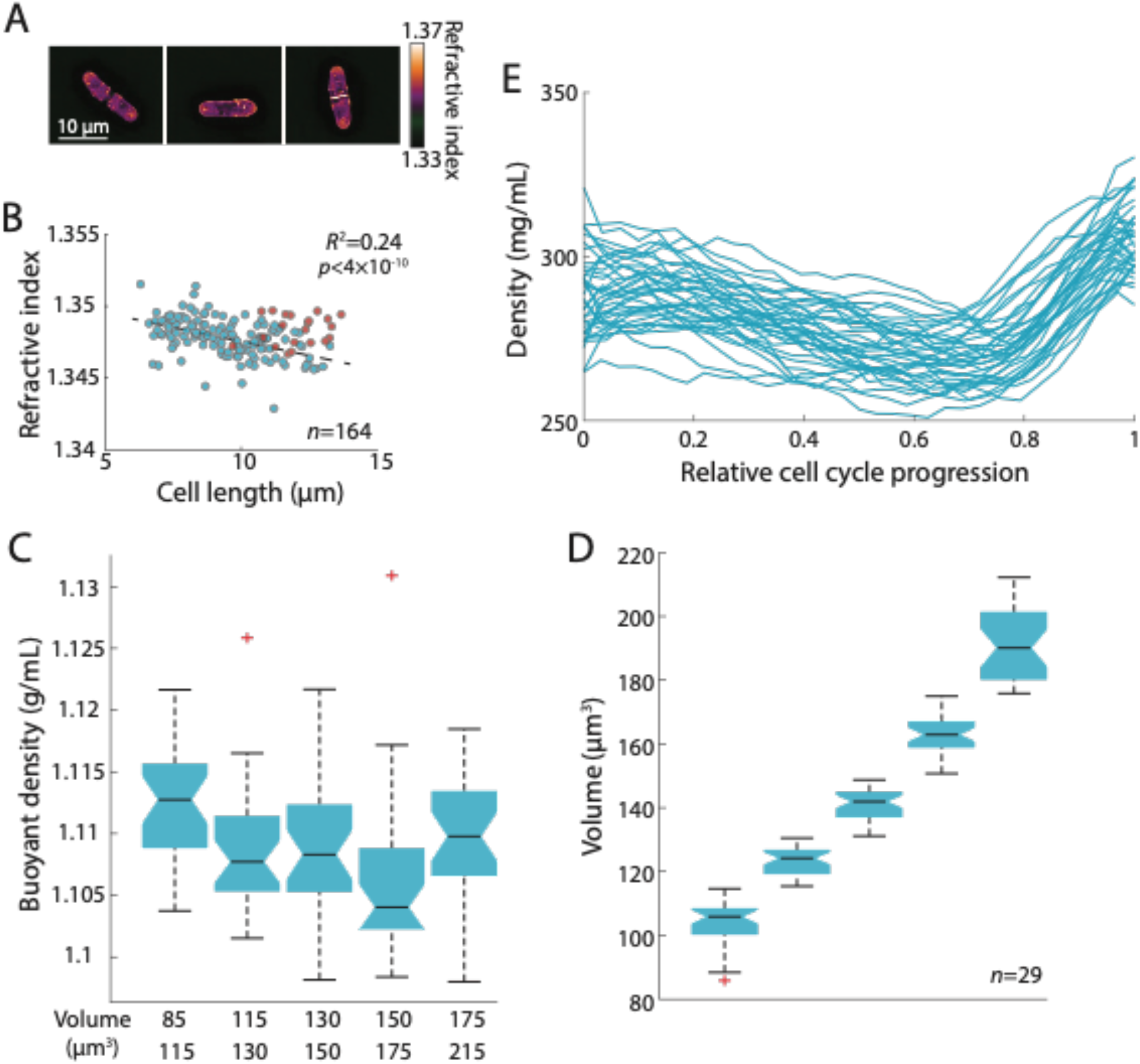
Measurements of buoyant density and buoyant mass using SMR and refractive index measurements using holography and QPI show cell cycle-dependent density variations. A) Representative holographic images of cells at an early, middle, and late stage in the cell cycle. B) Mean refractive index was calculated from holographic images of non-septated cells (blue) and septated cells (red). The negative correlation (Pearson’s correlation coefficient) of refractive index with non-septated cells (dashed line) indicates that refractive index decreases with increasing cell length. C) Replicate SMR experiment for data in Figure 2D. Medium-sized cells exhibited lower buoyant density than small and large cells, similar to QPI measurements (Figure 2C). Boxes indicate 25^th^ and 75^th^ percentiles and horizontal line indicates the median. Whiskers indicate the most extreme datapoints not considered outliers. D) Distribution of the volumes of cells in each bin of the buoyant density measurements in (C). Each bin contains 29 cells sorted by volume. E) QPI-based dry-mass density measurements of individual cells showed similar variation throughout the cell cycle as the population (Figure 2C).

**Supplemental Figure 3:**
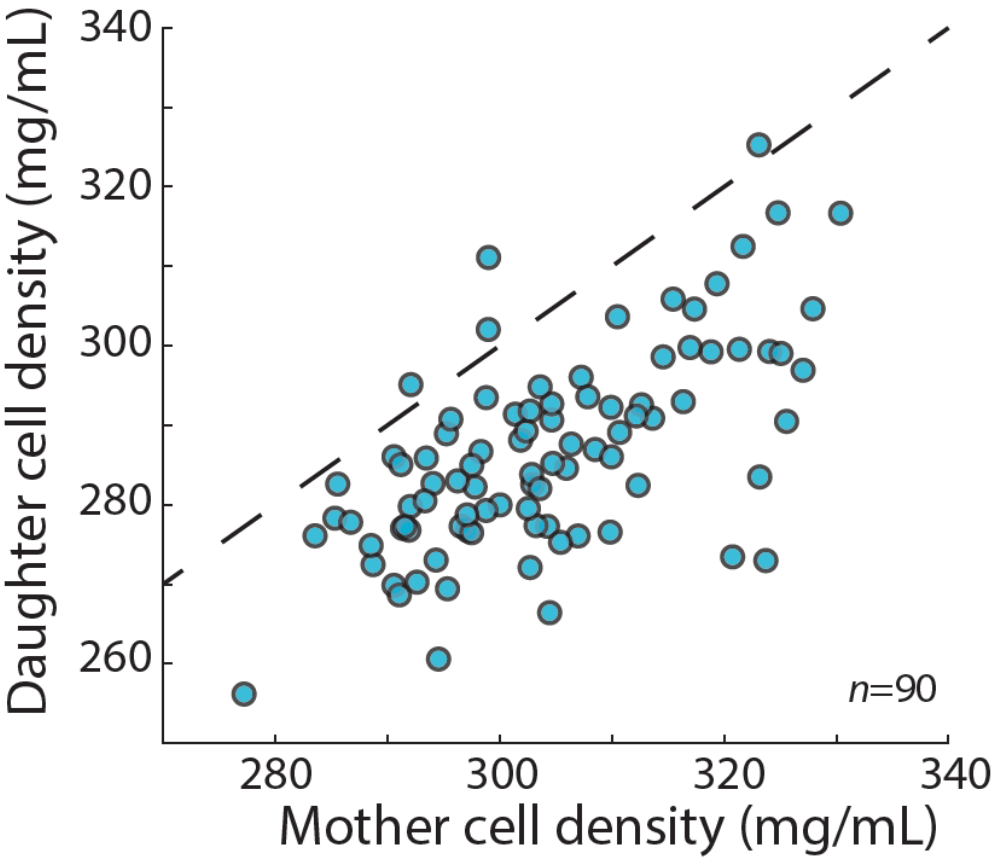
The mean density of *S. pombe* daughter cells was typically lower than the density of the mother cell. The densities of the mother cell and the associated daughter cells were measured from consecutive QPI density maps (5 min apart) directly before and after cell division, respectively. The daughter cell densities were then averaged. In large majority of cases, the newly born daughter cells were less dense than the mother cell 5 min beforehand. Dashed line indicates *y*=*x*.

**Supplemental Figure 4:**
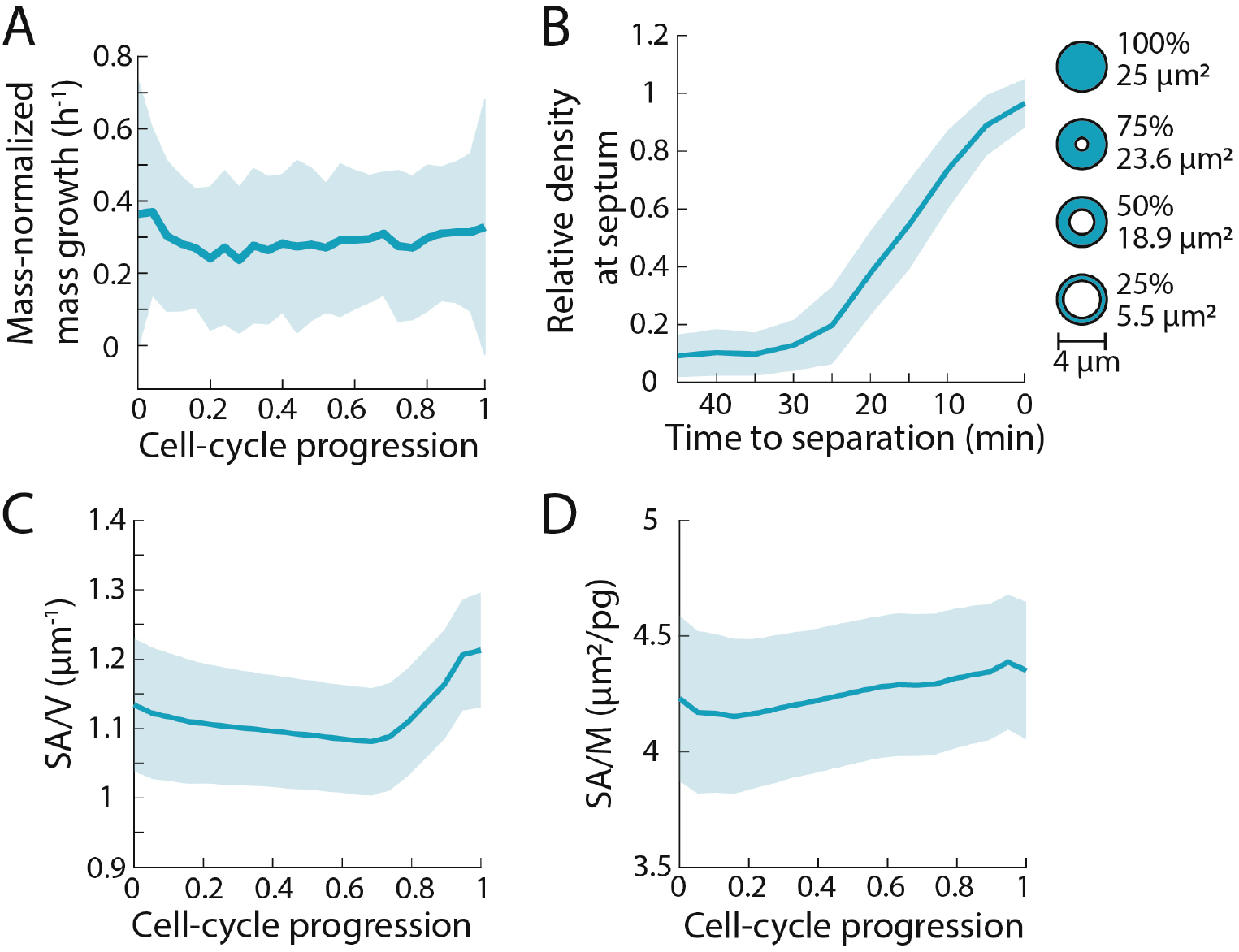
Cellular surface area-to-mass ratio varies less than dry-mass density during the *S. pombe* cell cycle. A) Mass increases exponentially throughout the cell cycle, as evidenced by the constant rate of normalized growth (1/*M dM*/*dt*). The dry mass of individual cells was measured as in Figure 2. Larger cells added more mass per unit time, a characteristic of exponential mass growth. Curves are mean values and shaded regions represent 1 standard deviation (*n*=78 cells). B) To measure surface area, we took into account the surface area added at the septum during cytokinesis. Septum growth was estimated by measurement of density at the septal region prior to division. The surface area of the septum was calculated by assuming a double-layered structure with a diameter of 4 μm. The local density was used to estimate the diameter of the opening, such that at 50% of maximal density, 50% of the cross-sectional diameter was assumed to be filled with septal wall. Curves are mean values and shaded regions represent 1 standard deviation (*n*=78 cells). C) Surface area to volume (SA/V) ratio increased at the end of the cell cycle, as volume growth slowed and surface area increased due to septum formation. Curves are mean values and shaded regions represent 1 standard deviation (*n*=78 cells). D) Surface area to mass (SA/M) ratio varied by only ~5%. Curves are mean values and shaded regions represent 1 standard deviation (*n*=78 cells). Thus, SA/M shows less variation than the ratio of volume to mass (the inverse of density).

**Supplemental Figure 5:**
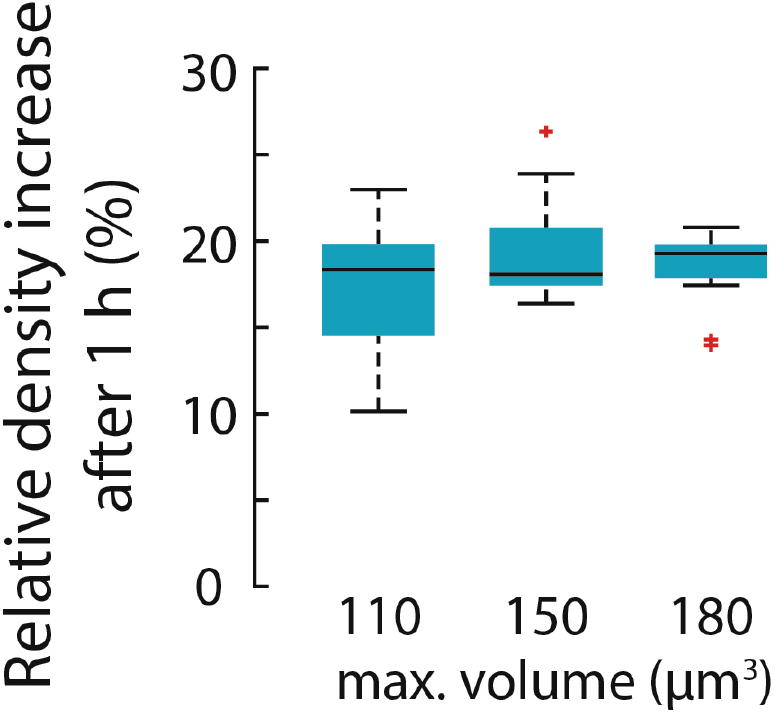
The increase in intracellular density due to treatment with the actin inhibitor latrunculin A was not dependent on cell size. Wild-type *S. pombe* cells were treated with 0.2 mM latrunculin A for 1 h and density maps were measured using QPI as in Figure 6. Density increase over 1 h normalized to mean density at the first time point was approximately invariant across initial cell sizes. Cells per bin, left to right: 10, 10, 12. Boxes indicate 25^th^ and 75^th^ percentiles and horizontal line indicates the median. Whiskers indicate the most extreme datapoints not considered outliers.

**Supplemental Figure 6:**
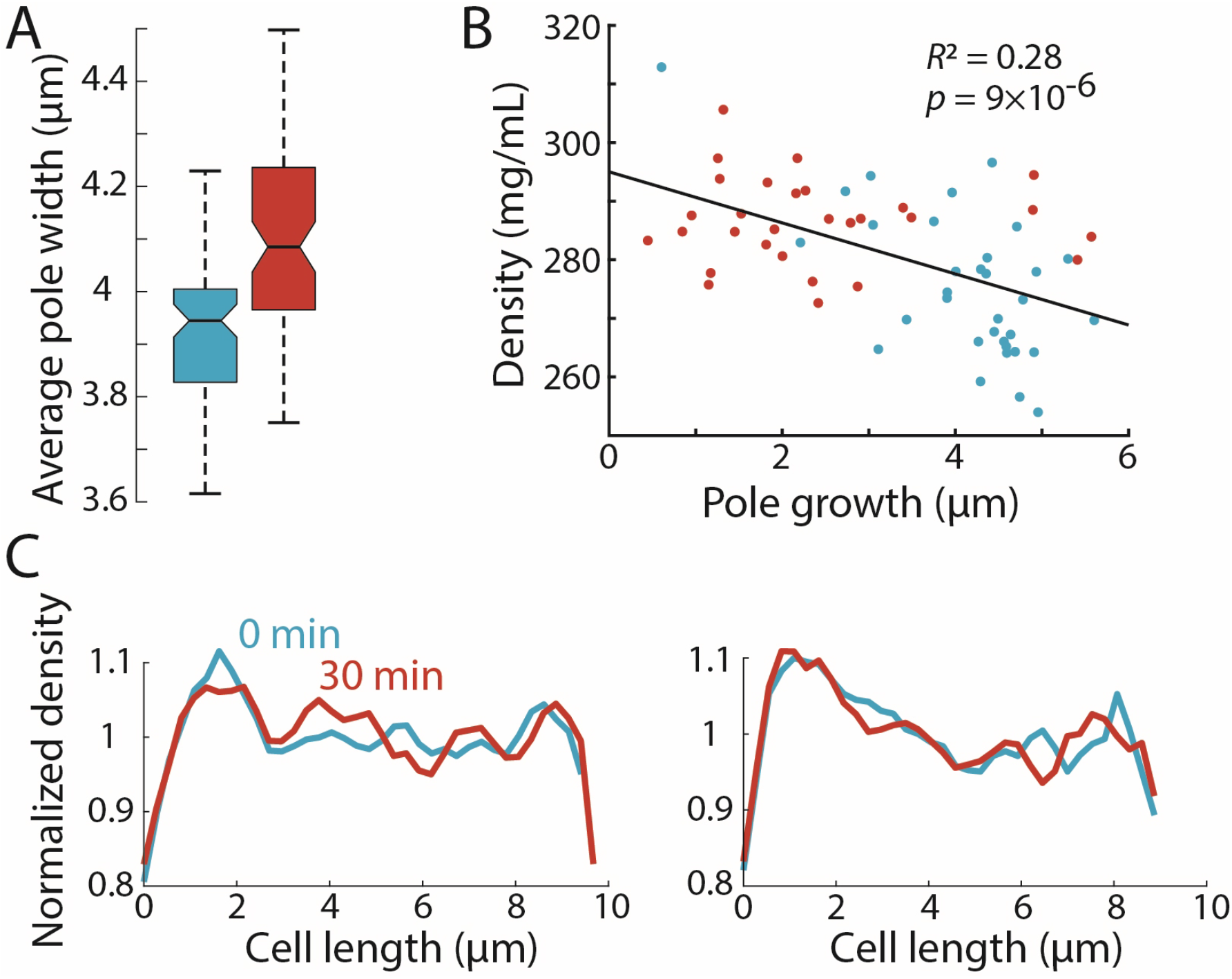
Spatial intracellular density gradients are present in cells with similar widths, and are stably maintained in cells treated with latrunculin A. A) The new cell end is wider than the old end. The mean width of the region 1.5-3 μm away from the cell pole was extracted for the new and old ends over time and averaged. The average old end width was ~0.15 μm smaller than that of new ends. Boxes indicate 25^th^ and 75^th^ percentiles and horizontal line indicates the median. B) To address a caveat that differences in QPI density maps may arise from differences in cellular height, we restricted our measurements of average pole density to poles with similar widths (between 3.9 and 4.1 μm, corresponding to the medians in (A)). Tip growth through the cell cycle and average density at the ends were plotted as in Figure 3C. The negative correlation (Pearson’s correlation coefficient) between pole growth and density observed in Figure 3B persisted, suggesting that the difference in measured density between poles is not due to differences in sample height. C) The density gradient was stable in latrunculin A-treated cells. Cells were treated with latrunculin A and imaged using QPI as in Figure 5. Normalized QPI density plots along the long axis of two individual cells before and 30 min after treatment with latrunculin A. The density gradient was maintained in the absence of F-actin and tip growth.

**Supplemental Figure 7:**
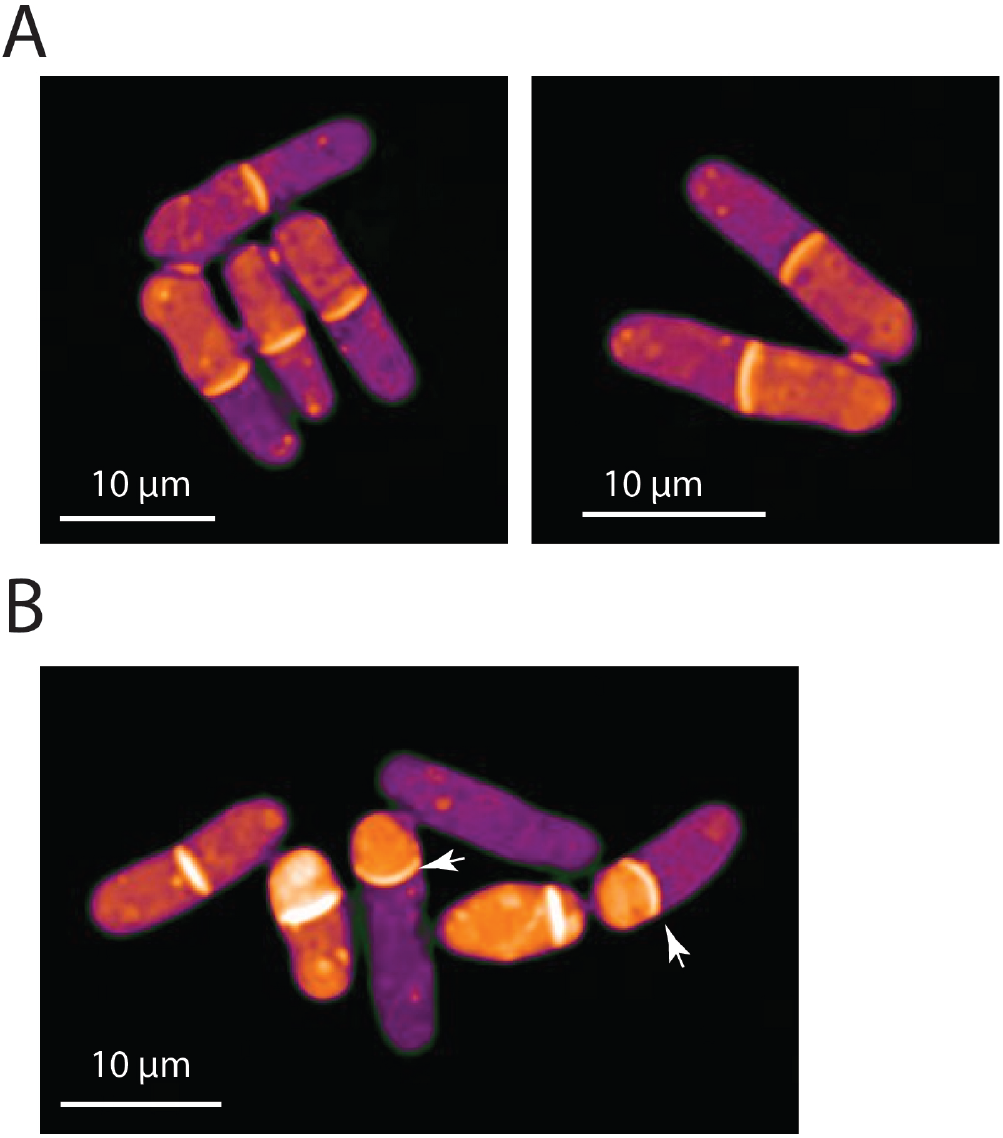
Septa bend away from the compartment of higher density in *mid2* and *cdc16* mutant cells. A) QPI density maps of *mid2Δ* cells delayed in cell separation often exhibited differences in density between sister-cell compartments, and in most cases the septum was bent away from the compartment of higher density. B) QPI density map of *cdc16* cells grown at 25 °C and then imaged on a microscope pre-heated to 34 °C; image shows *t*=3 h time point. These cells often exhibited an asymmetrically positioned septum, which formed a small cellular compartment that did not grow much and increased in density and septum bending (arrows).

## Notes

### Competing Interest Statement

The authors have declared no competing interest.

### Summary of Updates

Addition of new data on buoyant density measurements(Figure 2D, S2C,D), new data on density tracks of individual cells in Figure S2E, revision of author list, and revisions in text throughout.

## References

1. J. van den Berg, A. J. Boersma, B. Poolman, Microorganisms maintain crowding homeostasis. Nat Rev Microbiol 15, 309–318 (2017).

2. T. J. Mitchison, Colloid osmotic parameterization and measurement of subcellular crowding. Mol Biol Cell 30, 173–180 (2019).

3. G. E. Neurohr, A. Amon, Relevance and Regulation of Cell Density. Trends Cell Biol 30, 213–225 (2020).

4. H. X. Zhou, G. Rivas, A. P. Minton, Macromolecular crowding and confinement: biochemical, biophysical, and potential physiological consequences. Annu Rev Biophys 37, 375–397 (2008).

5. S. Oh et al., In situ measurement of absolute concentrations by Normalized Raman Imaging. bioRxiv 10.1101/629543, 629543 (2019).

6. S. Son et al., Resonant microchannel volume and mass measurements show that suspended cells swell during mitosis. J Cell Biol 211, 757–763 (2015).

7. E. Zlotek-Zlotkiewicz, S. Monnier, G. Cappello, M. Le Berre, M. Piel, Optical volume and mass measurements show that mammalian cells swell during mitosis. J Cell Biol 211, 765–774 (2015).

8. E. R. Rojas, K. C. Huang, Regulation of microbial growth by turgor pressure. Curr Opin Microbiol 42, 62–70 (2018).

9. G. E. Neurohr et al., Excessive Cell Growth Causes Cytoplasm Dilution And Contributes to Senescence. Cell 176, 1083–1097.e1018 (2019).

10. B. D. Knapp et al., Decoupling of Rates of Protein Synthesis from Cell Expansion Leads to Supergrowth. Cell Syst 9, 434–445.e436 (2019).

11. T. A. Zangle, M. A. Teitell, Live-cell mass profiling: an emerging approach in quantitative biophysics. Nat Methods 11, 1221–1228 (2014).

12. T. P. Burg et al., Weighing of biomolecules, single cells and single nanoparticles in fluid. Nature 446, 1066–1069 (2007).

13. Y. Park, C. Depeursinge, G. Popescu, Quantitative phase imaging in biomedicine. Nature Photonics 12, 578–589 (2018).

14. K. Lee et al., Quantitative phase imaging techniques for the study of cell pathophysiology: from principles to applications. Sensors (Basel, Switzerland) 13, 4170–4191 (2013).

15. E. Bostan, E. Froustey, M. Nilchian, D. Sage, M. Unser, Variational Phase Imaging Using the Transport-of-Intensity Equation. IEEE Trans Image Process 25, 807–817 (2016).

16. J. M. Mitchison, P. Nurse, Growth in cell length in the fission yeast Schizosaccharomyces pombe. J Cell Sci 75, 357–376 (1985).

17. B. Rappaz et al., Noninvasive characterization of the fission yeast cell cycle by monitoring dry mass with digital holographic microscopy. J Biomed Opt 14, 034049 (2009).

18. W. H. Grover et al., Measuring single-cell density. Proc Natl Acad Sci U S A 108, 10992–10996 (2011).

19. F. Feijó Delgado et al., Intracellular water exchange for measuring the dry mass, water mass and changes in chemical composition of living cells. PloS one 8, e67590–e67590 (2013).

20. A. B. Martín-Cuadrado, E. Dueñas, M. Sipiczki, C. R. Vázquez de Aldana, F. del Rey, The endo-beta-1,3-glucanase eng1p is required for dissolution of the primary septum during cell separation in Schizosaccharomyces pombe. J Cell Sci 116, 1689–1698 (2003).

21. B. S. Hercyk, U. N. Onwubiko, M. E. Das, Coordinating septum formation and the actomyosin ring during cytokinesis in Schizosaccharomyces pombe. Mol Microbiol 112, 1645–1657 (2019).

22. E. Atilgan, V. Magidson, A. Khodjakov, F. Chang, Morphogenesis of the Fission Yeast Cell through Cell Wall Expansion. Curr Biol 25, 2150–2157 (2015).

23. V. Simanis, Pombe’s thirteen – control of fission yeast cell division by the septation initiation network. Journal of Cell Science 128, 1465 (2015).

24. S. Ray et al., The mitosis-to-interphase transition is coordinated by cross talk between the SIN and MOR pathways in Schizosaccharomyces pombe. Journal of Cell Biology 190, 793–805 (2010).

25. S. G. Martin, R. A. Arkowitz, Cell polarization in budding and fission yeasts. FEMS Microbiology Reviews 38, 228–253 (2014).

26. A. Papagiannakis, B. Niebel, E. C. Wit, M. Heinemann, Autonomous Metabolic Oscillations Robustly Gate the Early and Late Cell Cycle. Mol Cell 65, 285–295 (2017).

27. B. Novak, J. M. Mitchison, Change in the rate of CO2 production in synchronous cultures of the fission yeast Schizosaccharomyces pombe: a periodic cell cycle event that persists after the DNA-division cycle has been blocked. J Cell Sci 86, 191–206 (1986).

28. J. M. Mitchison, “Growth During the Cell Cycle” in International Review of Cytology. (Academic Press, 2003), vol. 226, pp. 165–258.

29. P. Nurse, P. Thuriaux, K. Nasmyth, Genetic control of the cell division cycle in the fission yeast Schizosaccharomyces pombe. Molecular and General Genetics MGG 146, 167–178 (1976).

30. I. Hagan, M. Yanagida, Novel potential mitotic motor protein encoded by the fission yeast cut7+ gene. Nature 347, 563–566 (1990).

31. M. Minet, P. Nurse, P. Thuriaux, J. M. Mitchison, Uncontrolled septation in a cell division cycle mutant of the fission yeast Schizosaccharomyces pombe. J Bacteriol 137, 440–446 (1979).

32. K. Z. Pan, T. E. Saunders, I. Flor-Parra, M. Howard, F. Chang, Cortical regulation of cell size by a sizer cdr2p. eLife 3, e02040 (2014).

33. D. R. Mutavchiev, M. Leda, K. E. Sawin, Remodeling of the Fission Yeast Cdc42 Cell-Polarity Module via the Sty1 p38 Stress-Activated Protein Kinase Pathway. Current biology : CB 26, 2921–2928 (2016).

34. F. Chang, S. G. Martin, Shaping fission yeast with microtubules. Cold Spring Harb Perspect Biol 1, a001347 (2009).

35. J. Muñoz et al., Extracellular cell wall β(1,3)glucan is required to couple septation to actomyosin ring contraction. The Journal of cell biology 203, 265–282 (2013).

36. A. Berlin, A. Paoletti, F. Chang, Mid2p stabilizes septin rings during cytokinesis in fission yeast. J Cell Biol 160, 1083–1092 (2003).

37. A. K. Bryan, A. Goranov, A. Amon, S. R. Manalis, Measurement of mass, density, and volume during the cell cycle of yeast. Proc Natl Acad Sci U S A 107, 999–1004 (2010).

38. J. M. Mitchison, The growth of single cells. I. Schizosaccharomyces pombe. Exp Cell Res 13, 244–262 (1957).

39. V. Stonyte, E. Boye, B. Grallert, Regulation of global translation during the cell cycle. J Cell Sci 131(2018).

40. M. Godin et al., Using buoyant mass to measure the growth of single cells. Nat Methods 7, 387–390 (2010).

41. X. Liu, S. Oh, L. Peshkin, M. W. Kirschner, Computationally Enhanced Quantitative Phase Microscopy Reveals Autonomous Oscillations in Mammalian Cell Growth. bioRxiv 10.1101/631119, 631119 (2019).

42. T. P. Miettinen, J. H. Kang, L. F. Yang, S. R. Manalis, Mammalian cell growth dynamics in mitosis. Elife 8(2019).

43. E. R. Oldewurtel, Y. Kitahara, B. Cordier, G. Özbaykal, S. van Teeffelen, Bacteria control cell volume by coupling cell-surface expansion to dry-mass growth. bioRxiv 10.1101/769786, 769786 (2019).

44. D. Y. Parkinson, G. McDermott, L. D. Etkin, M. A. Le Gros, C. A. Larabell, Quantitative 3-D imaging of eukaryotic cells using soft X-ray tomography. J Struct Biol 162, 380–386 (2008).

45. K. Kume et al., A systematic genomic screen implicates nucleocytoplasmic transport and membrane growth in nuclear size control. PLoS Genet 13, e1006767 (2017).

46. N. Erjavec, M. Cvijovic, E. Klipp, T. Nyström, Selective benefits of damage partitioning in unicellular systems and its effects on aging. Proc Natl Acad Sci U S A 105, 18764–18769 (2008).

47. W. Choi et al., Tomographic phase microscopy. Nat Methods 4, 717–719 (2007).

48. O. Tolde et al., Quantitative phase imaging unravels new insight into dynamics of mesenchymal and amoeboid cancer cell invasion. Sci Rep 8, 12020 (2018).

49. C. V. Harding, C. Feldherr, Semipermeability of the nuclear membrane in the intact cell. J Gen Physiol 42, 1155–1165 (1959).

50. S. Moreno, A. Klar, P. Nurse, Molecular genetic analysis of fission yeast Schizosaccharomyces pombe. Methods Enzymol 194, 795–823 (1991).

51. A. D. Edelstein et al., Advanced methods of microscope control using μManager software. J Biol Methods 1(2014).

52. T. Ursell et al., Rapid, precise quantification of bacterial cellular dimensions across a genomic-scale knockout library. BMC Biol 15, 17 (2017).

53. A. Meyers et al., Lipid Droplets Form from Distinct Regions of the Cell in the Fission Yeast Schizosaccharomyces pombe. Traffic 17, 657–669 (2016).

54. J. H. Kang et al., Noninvasive monitoring of single-cell mechanics by acoustic scattering. Nat Methods 16, 263–269 (2019).

